# Temperature response of wheat affects final height and the timing of stem elongation under field conditions

**DOI:** 10.1101/756700

**Authors:** Lukas Kronenberg, Steven Yates, Martin P. Boer, Norbert Kirchgessner, Achim Walter, Andreas Hund

**Affiliations:** Crop Science, Institute of Agricultural Sciences, ETH Zürich, 8092 Zurich, Switzerland; Molecular Plant Breeding, Institute of Agricultural Sciences, ETH Zürich, 8092 Zurich, Switzerland; Biometris, Wageningen University & Research, 6708 PB Wageningen, The Netherlands

**Keywords:** development, field phenotyping, GWAS, LIDAR, physiology, plant height, temperature response, wheat

## Abstract

In wheat, temperature affects the timing and intensity of stem elongation (SE). Genetic variation for this process is therefore important for adaptation. This study investigates the genetic response to temperature fluctuations during SE and its relationship to phenology and height. Canopy height of 315 wheat genotypes (GABI wheat panel) was scanned twice weekly in the field phenotyping platform (FIP) of ETH Zurich using a LIDAR. Temperature response was modelled using linear regressions between SE and mean temperature in each measurement interval. This led to a temperature–responsive (slope) and a temperature-irresponsive (intercept) component.

The temperature response was highly heritable (H^2^ = 0.81) and positively related to a later start and end of SE as well as final height. Genome-wide association mapping revealed three temperature-responsive and four temperature-irresponsive quantitative trait loci (QTL). Furthermore, putative candidate genes for temperature-response QTL were frequently related to the flowering pathway in *A. thaliana*, whereas temperature-irresponsive QTLs corresponded with growth and reduced height genes. In combination with *Rht* and *Ppd* alleles, these loci, together with the loci for the timing of SE accounted for 71% of the variability in height.

This demonstrates how high-throughput field phenotyping combined with environmental covariates can contribute to a smarter selection of climate-resilient crops.

**Highlight:** We measured ambient temperature response of stem elongation in wheat grown under field conditions. The results indicate that temperature response is highly heritable and linked to the flowering pathway.

## Introduction

Temperature is a major abiotic factor affecting plant growth and development. As a consequence of global warming, wheat production could decrease by 6% per °C global temperature increase (Asseng *et al*., 2015). While heat stress during critical stages can drastically reduce yield (Gibson and Paulsen, 1999; Farooq *et al*., 2011), warm temperatures can decrease yield by accelerating development and thereby shortening critical periods for yield formation (Fischer, 1985; Slafer and Rawson, 1994). Despite the clear effect of temperature on growth and phenology, little is known about the genotype-specific response pattern to varying temperature conditions during crop development and its genetic control. We therefore aimed to quantify the genotype-specific temperature-responsiveness of European winter wheat during the stem elongation phase.

Stem elongation (SE) is a critical phase for yield formation in wheat. It occurs between the phenological stages of terminal spikelet initiation and anthesis (Slafer *et al*., 2015). The start of SE coincides with the transition from vegetative to reproductive development, when the apex meristem differentiates from producing leaf primordia to producing spikelet primordia (Trevaskis *et al*., 2007; Kamran *et al*., 2014). During SE, florets are initiated at the spikelets until booting (Kirby, 1988; Slafer *et al*., 2015). An increased duration of SE increases the number of fertile florets due to longer spike growth and higher dry matter partitioning to the spike (González *et al*., 2003). This in turn increases the number of grains per spike and therefore yield (Fischer, 1985). Modifying the timing of the critical phenological stages (transition to early reproductive phase and flowering), and thus SE duration, has been proposed as a way to increase wheat yield (Slafer *et al*., 1996; Miralles and Slafer, 2007; Whitechurch *et al*., 2007) or at least to mitigate adverse climate change effects on yield e.g. by enhancing earliness to escape heat during flowering (Chapman *et al*., 2012; Hernandez-Ochoa *et al*., 2019). The recent warming trend causes a faster advancement in phenology. For example flowering time occurred earlier in Germany throughout the past decade, which is attributable to both, increased temperature and selection for early flowering (Rezaei *et al*., 2018).

Final height is also an important yield determinant. During the “green revolution” wheat yields increased by the introduction of reduced height genes (*Rht*). The resulting dwarf and semi dwarf varieties benefit from improved resource allocation from the stem to the spike and reduced lodging, allowing more intensive nitrogen application (Hedden, 2003). Gibberellin insensitive *Rht* genes (*Rht-A1, Rht-B1*, and *Rht-D1*) were shown to limit cell wall extensibility which decreases growth rates (Keyes *et al*., 1989) without affecting development (Youssefian *et al*., 1992). Moreover, the allele *Rht-B1c* (Wu *et al*., 2011) and the GA sensitive *Rht12* dwarfing gene (Chen *et al*., 2013) delay heading.

The main abiotic factors affecting the timing of floral initiation and flowering are temperature and photoperiod; with temperature affecting both vernalisation and general rate of development (Slafer *et al*., 2015). These developmental transitions are controlled by major genes involved in the flowering pathway; namely: vernalisation (*Vrn)*, photoperiod (*Ppd*) and earliness per se (*Eps*) genes (Slafer *et al*., 2015). The *Ppd* and *Vrn* genes define photoperiod and vernalisation requirements which jointly enable the transition to generative development and define time to flowering. Whereas *Eps* genes fine tune the timing of floral transition and flowering, after vernalisation and photoperiod requirements are fulfilled (Zikhali and Griffiths, 2015). While vernalisation and photoperiod response are well known, the role of temperature *per se* remains less clear. Temperature affects all developmental phases and warmer ambient temperatures generally accelerate growth and development in crops (Slafer and Rawson, 1994, 1995a,c; Atkinson and Porter, 1996; Fischer, 2011; Slafer *et al*., 2015). But it is unclear, if temperature response governs growth rate and development independently. If so, the question remains, whether there is enough genetic variability in temperature response to be used in a breeding context (Parent and Tardieu, 2012).

Genotypic variation for growth response to temperature was reported for wheat leaf elongation rate (Nagelmüller *et al*., 2016), as well as for canopy cover growth (Grieder *et al*., 2015). Kiss et al. (2017) reported significant genotype by temperature interactions in the timing of SE as well as temperature dependent differences in the expression of *Vrn* and *Ppd* genes under controlled conditions. Under field conditions, the response of stem elongation to temperature has not yet been investigated in high temporal resolution.

In recent years, new high throughput phenotyping technologies have enabled monitoring plant height with high accuracy and frequency in the field (Bendig *et al*., 2013; Friedli *et al*., 2016; Holman *et al*., 2016; Aasen and Bareth, 2018; Hund *et al*., 2019). We have previously demonstrated that the ETH field phenotyping platform (FIP, Kirchgessner *et al*., 2016) can be used to accurately track the development of canopy height in a large set of wheat genotypes using terrestrial laser scanning (Kronenberg *et al*., 2017). Considerable genotypic variation was detected for the start and end of SE which correlated positively with final canopy height (Kronenberg *et al*., 2017).

While many temperature-independent factors affecting plant height are known, the influences of temperature-dependent elongation and timing of the elongation phase is less clear. We hypothesise that apart from temperature-independent factors, there is a genotype specific response to ambient temperature which affects growth as well as the timing of developmental stages. To address this, we aimed to dissect the process towards final height into the following components: i) temperature-independent elongation, ii) temperature-dependent elongation and iii) the duration of the elongation phase determined by the start and end of the process. To achieve this we present a method to assess and measure these three processes under field conditions by means of high-frequency, high-throughput phenotyping of canopy height development. The resulting data were combined with genetic markers to identify quantitative trait loci controlling the aforementioned processes.

## Material and Methods

### Experimental setup, phenotyping procedures and extracted traits

Field experiments were conducted in the field phenotyping platform FIP at the ETH research station in Lindau-Eschikon, Switzerland (47.449°N, 8.682°E, 520 m a.s.l.; soil type: eutric cambisol). We used a set of approximately 330 winter wheat genotypes (335 – 352 depending on the experiment) comprising current European elite cultivars (GABI Wheat; Kollers *et al*., 2013), supplemented with thirty Swiss varieties. These were monitored over three growing seasons in 2015, 2016 and 2017. Briefly, the field experiments were conducted in an augmented design with two replications per genotype using micro plots with a size of 1.4 by 1.1 m. In the growing season 2017, the experiment was repeated again, with minor changes in genotypic composition. This resulted in 328 genotypes present across all three experiments. Details about the experimental setup for the growing seasons 2015 and 2016 are described in Kronenberg et al. (2017).

Canopy height was measured twice weekly from the beginning of shooting (BBCH 31) using a light detection and ranging (LIDAR) scanner (FARO R Focus3D S 120; Faro Technologies Inc., Lake Mary USA) mounted on the FIP (Kirchgessner *et al*., 2016). Height measurements were concluded when no further height increase was observed in any of the genotypes. Canopy height data was extracted from the LIDAR data as described in Kronenberg et al. (2017).

The start, end, and duration of SE as well as final canopy height (FH) were extracted from the height data following Kronenberg et al. (2017): FH was defined as the point, after which no increase in height was observed in several consecutive measurements. Normalized canopy height was calculated as percent of FH at each day of measurement for every plot and then linearly interpolated between measurement points. Growing degree-days after sowing until 15% FH (GDD_15_) and 95% FH (GDD_95_) were used as proxy traits for start and end of SE, respectively. SE duration was recorded in thermal time (GDD_SE_) as well as in calendar days (time_SE_), as the difference between GDD_95_ and GDD_15_ (Kronenberg *et al*., 2017). Heading date was recorded as GDD (heading_GDD_) when 50% of the spikes were fully emerged from the flag leaf sheath (BBCH 59; Lancashire *et al*., 1991). Heading data for 2015 could not be evaluated due to insufficient data availability. Therefore, a third year of heading data was gathered in 2018, when the experiment was repeated again as described in Anderegg et al., (2020).

In order to investigate short-term growth response to temperature, average daily stem elongation rates (SER) were calculated for each plot as the difference (Δ) in canopy height (CH) between consecultive time points (t):

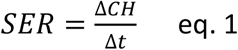

### Extracting growth response to temperature

Temperature response was modelled by regressing average daily SER against average temperature of the respective interval for each plot within the respective year following

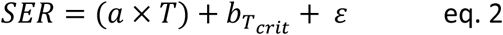

where *T* is the ambient temperature, *a* is the coefficient of the linear regression (*i.e*. growth response to ambient temperature; slp_SER^~^T_) and *ε* denotes the residual error. *b_Tcrit_* is the model intercept at the temperature, at which the correlation between intercept (int_SER^~^T_) and slope is zero (*see below*). Per definition, the intercept of a linear model would be calculated at T = 0 °C, *i.e*. far outside the range of observed temperatures. In the observed data, the intercept at T = 0°C correlated strongly negative with the slope (Fig. 1A) and thus, did not add much additional information concerning the performance of the evaluated genotypes. Likewise, an intercept at 20°C, at the upper range of the observed data correlated strongly positive with the slope (Fig 1A). Grieder *et al*. (2015) performed a similar analysis for the canopy cover development during winter and found a similar, strongly negative correlation between temperature response (slope) and growth at 0°C (intercept). We sequentially calculated the intercept at 0.01°C increments between 1°C and 22°C for each plot within a year. Subsequently, we calculated the Pearson correlation coefficient between the common slope of these models and each of the different intercepts (Fig 1A). Based on this sequence, we empirically determined the critical temperature value (*T_crit_*) at which the correlation between slope and intercept was zero (Fig 1A). Hence, *T_crit_* is defined as the temperature at which intercept and slope are independent. Due to this independence, the value of the intercept at *T_crit_* can be interpreted as intrinsic growth component independent of temperature response herein referred to as “vigour”. Following this, two genotypes can show the same vigour but differ markedly in temperature response (Fig. 1B), have the same temperature response but differ in vigour (Fig. 1C), or differ for both, temperature response and vigour (Fig. 1D).

**Fig. 1:**
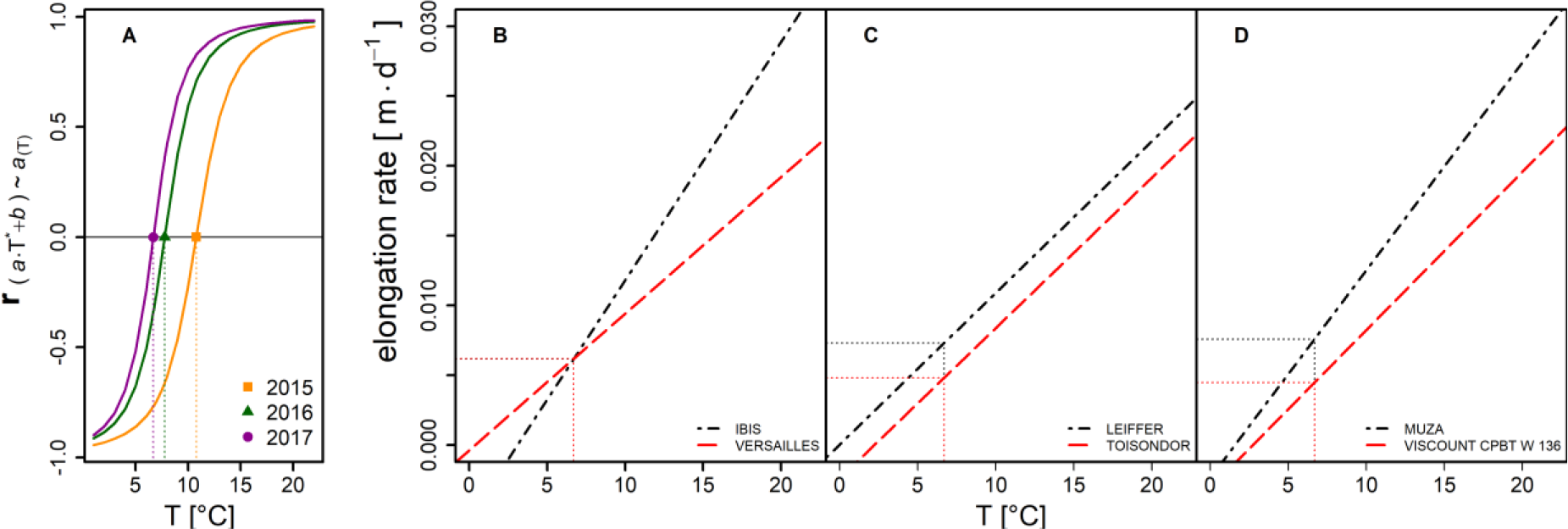
Illustration and interpretation for the parameters of the applied temperature response model (eq. 2). **A:** Distribution of Pearson correlation coefficients between intercept and slope of the linear model for individual years, depending on the temperature, at which the intercept is calculated. Dotted vertical lines indicate the critical temperature (T_crit_) for individual years used to calculate the intercept. **B-D**: Illustration of the relation between intercept and slope on contrasting genotypes (dashed and dash-dotted lines). **B**: same vigour but different in temperature response. **C** both have the same T-response but differ in vigour. **D** Genotypes differ in vigour as well as in T-response. Horizontal and vertical dotted lines indicate vigour and T_crit_ respectively. The two contrasting genotypes per example (B-D) were selected from the 2017 data based on their vales for slope and intercept.

### Statistical Analysis

The data were processed stepwise as follows: i) correction for design factors and spatial trends, ii) application of linear models to each plot to determine growth-response to temperature and iii) prediction of adjusted means and calculation of the heritability for all traits across years.

The spatial correction of canopy height per measurement time point was done using the R-package SpATS (Rodríguez-Álvarez *et al*., 2018) following

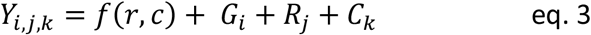

where *Y* is the phenotypic value for the a plot in the j^th^ row in the k^th^ column planted with the i^th^ genotype, *f(r,c)* is a smoothed bivariate surface defined over rows and ranges of a virtual grid, *G_i_* is the effect of the i^th^ genotype (i = 1,…, n; n= 335 – 352 depending on the year), *R_j_* is the effect of the j^th^ row in the virtual grid and *C_k_* is the effect of the k^th^ range. With the number of genotypes, the number of ranges/rows varied across years (17/21 in 2015, 18/20 in 2016 and 18/21 in 2017, respectively). The replications were arranged diagonally in this grid with a gap of five rows and ranges between them. Thus, e.g. in 2017, replication 1 was at rows 1-21 and ranges 1-18; replication 2 was at rows 24-41 and ranges 27-47 of the virtual grid.

The function *f(r,c)* describing the bivariate surface can be decompose in a nested-type ANOVA structure as describe by Rodríguez-Álvarez et al. (2018). The number of spline points was set to 2/3 of the total number of rows and ranges in the virtual grid, respectively.

From this model the predicted genotypic best linear unbiased estimates (BLUEs) plus residual error were kept as spatially corrected plot value. Thus, these new plot values were corrected for spatial effect as well as for the random row and range effects and used for the subsequent dynamic model. For a visualisation of the applied SpATS correction, see Fig. S1.

Genotypic BLUEs across the three seasons were calculated for all traits using the R-package asreml-4 (Butler, 2018) following

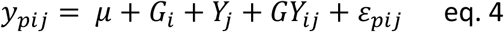

where *y_pij_* is the spatialy corrected plot value of the respective trait (FH, GDD_15_, GDD_95_, GDD_SE_, time_SE_, int_SER^~^T_ or slp_SER^~^T_), μ is the overall mean, G_i_ the fixed effect of the genotypes common in all three years (i = 1,…,328), *Y*_j_ is the fixed effect of the year (j = 2015,…,2017), *G_i_Y_j_* ist the random genotype-by-year interaction and *ε_pij_* is the residual error.

In order to estimate best linear unbiased predictors and heritability (H^2^) across years, *G_i_* in eq. 3 was set as random term and heritability was calculated following (Falconer and Mackay, 1996) using

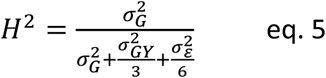

where *H^2^* is the broad sense heritability, 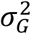 is the genotypic variance, 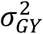 is the genotype-by-year interaction variance and 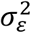 is the residual variance. For heading data, only one replicate per year was available. Plot corrected values were extracted using SpATS (Rodríguez-Álvarez *et al*., 2018) and heritability across three years was calculated by omitting the *GY* term in eq. 4 and dividing the residual variance by three, based on the three available year-site replications.

Genotypic BLUEs across three years were used for subsequent correlation analysis and genome wide association study (GWAS). All statistical analyses were performed in the R environment (R Core Team, 2015).In order to investigate the relationship between FH, temperature response and vigour and to test for confounding *Rht* or *Ppd* effects on temperature response, FH was modelled using the linear model

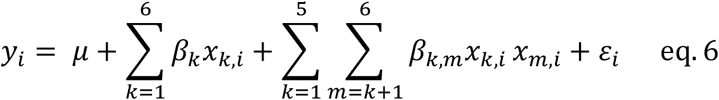

where *y_i_* is FH of the i^th^ genotype, *μ* is the model intercept, *β*_1−6_ are the main effect estimates of *x* = slp_SER^~^T_, int_SER^~^T_, GDD_SE_, *Rht-B1, Rht-D1* or *Ppd-D1*, respectively. *β*_1,2_ − *β*_5,6_ are all two-way interaction effects (n = 15) and *ε_i_* is the residual error. Genotypic data for *Rht-B1, Rht-D1* and *Ppd-D1* alleles was available for 301 genotypes obtained from Kollers et al. (2014). There, genotyping of the *Rht-1* alleles was performed using pcr markers (Ellis *et al*., 2002), while *Ppd-D1* alleles were genotyped by presence or absence of a 2-kb insertion using specific primers (Beales *et al*., 2007; Kollers *et al*., 2014)

### Association study

GWAS was performed on the different traits to compare the phenotypic correlations with the underlying genetic architecture of the traits. As a positive control FH data made in Germany and France by Zanke et al., (2014b) was also compared and analysed.

Genotyping data was made previously by the GABI wheat consortium represented by the Leibniz Institute of Plant Genetics and Crop Plant Research (IPK; Zanke *et al*., 2014a) using the 90K illumina SNP-chip (Cavanagh *et al*., 2013; Wang *et al*., 2014). Monomorphic SNPs were discarded. The remaining markers were mapped to the IWGSC reference genome (Consortium (IWGSC) *et al*., 2018) by BLASTN search using an E-value threshold < 1e^−30^. The genome position with the lowest E-value was assigned as the respective marker location. Markers that could not be unequivocally positioned were dropped. After filtering SNPs with a minor allele frequency and missing genotype rate < 0.05, a total of 13,450 SNP markers and 315 genotypes remained in the set. The reference genome position of *Rht, Ppd, Vrn* and putative *Eps* genes was determined with BLASTN search as described above using published GenBank sequences (Table S1).

To mitigate against multiple testing, relatedness and population structure; three different methods were used to calculate marker trait associations (MTA) between phenotypic BLUPs and SNP markers:

i. We used a mixed linear model (MLM) including principal components among marker alleles as fixed effects and kinship as random effect to account for population structure (Zhang *et al*., 2010). This approach was chosen to stringently prevent type I errors. The MLM GWAS was performed using the R Package GAPIT (v.2, Tang *et al*., 2016). Kinship was estimated according to VanRaden (2008).
ii. In a generalised linear model (GLM) framework implemented in PLINK (Purcell *et al*., 2007), association analysis was performed using SNP haplotype blocks consisting of adjacent SNP triplets. Using haplotype blocks takes the surrounding region of a given SNP into account, thus increasing the power to detect rare variants (Purcell *et al*., 2007)
iii. Finally, the FarmCPU method (Liu *et al*., 2016) was used, which is also implemented in GAPIT. FarmCPU tests individual markers with multiple associated markers as covariates in a fixed effect model. Associated markers are iteratively used in a random effect model to estimate kinship. Confounding between testing markers and kinship is thus removed while controlling type I error, leading to increased power (Liu *et al*., 2016).

For all methods, a Bonferroni correction was applied to the pointwise significance threshold of α = 0.05, to avoid false-positives. Hence, only markers above −log10(*P*-value) >= 5.43 considered significant.

Linkage disequilibrium (LD) among markers was estimated using the squared correlation coefficient (*r*^2^) calculated with the R package SNPrelate (Zheng *et al*., 2012). A threshold of *r*^2^ = 0.2 (Gaut and Long, 2003) was applied to calculate the chromosome specific distance threshold of LD decay. Putative candidate genes were identified by searching the IWGSC annotation of the reference genome (Consortium (IWGSC) *et al*., 2018) for genes associated with growth and development within the LD distance threshold around the respective MTA.

## Results

### Phenotypic results

We measured the canopy height of 710 – 756 plots per year, containing 335 – 352 wheat genotypes, for three consecutive years. In each season, measurements were made between 17 and 22 times during SE. Plot based growth rates within single years extracted from these data indicate a clear relation between growth and temperature for the period of SE, as depicted in Fig. 2. Towards the end of the measurement period in June, there was a larger deviation, which was also reflected in the quality of plot based linear model fits of SER versus temperature (*see* eq. 2), summarized in Fig. S2. For the 2015 and especially the 2016 experiment, R^2^ values were low and except for the 2017 experiment, the parameter estimates were not statistically significant (Fig. S2A). Inspection of the best and worst model fits however shows, that failure of fitting the model for single plots was levelled out by the replications within genotypes (Fig. S2B). The weak model fits did therefore not affect the genotype ranking of adjusted means across replications. Analysis of variance revealed significant (*P* < 0.001) genotypic effects for both slp_SER^~^T_ and int_SER^~^T_ across three years. Both traits showed high heritabilities across years (H^2^ = 0.81 for slp_SER^~^T_ and H^2^ = 0.77 for int_SER^~^T_; Table 1). Using the BLUEs of slp_SER^~^T_, int_SER^~^T_ and temperature sum for SE (GDD_SE_), FH could be predicted with high accuracy across different years (0.85 <= R^2^ <= 0.89) by training a linear model on the BLUEs of one year and predicting it on the BLUEs of another independent year. In order to account for possible confounding *Rht* and *Ppd* effects, the allelic status of *Rht-B1, Rht-D1* and *Ppd-D1* were included as contrasts in the model (Table 2). Of the 301 genotypes with available data 7 % and 58 % carried the dwarfing alleles *Rht-B1b* and *Rht-D1b*, respectively, and 13 % carried the photoperiod insensitive allele *Ppd-D1a*. Training the model on the 3-year BLUEs resulted in a prediction accuracy of single years between R^2^ = 0.94 and R^2^= 0.95 (Fig. 3). Type II ANOVA revealed significant effects for slp_SER^~^T_, int_SER^~^T_, GDD_SE_, *Rht-B1* and *Rht-D1*. A significant (*P* < 0.05) interaction effect was found between *Rht-D1* and *Ppd-D1*. Furthermore, weak interactions (*P* < 0.1) were found for *Rht-B1:Ppd-D1*, int*_SER^~^T_:Rht-B1* and int_SER^~^T_*:Rht-D1* (Table 2). High heritabilities across three years (0.54 <= H^2^ <= 0.98; Table 1), were also found for the other traits; FH, start of SE, end of SE, SE duration and heading. All traits showed moderate genotype by year interaction effects which were smaller (except for SE duration) than the genotypic effects across years (Table 1).

**Table 1:** Variance components and heritabilities for all investigated traits.

| Trait | var. comp. | estimate | std.error | z.ratio | % tot. var. | Heritability |
| --- | --- | --- | --- | --- | --- | --- |
| int<sub>SER~T</sub> | Gen_ID | 7.024E-07 | 7.219E-08 | 9.730 | 49.01 | 0.77 |
|  | Gen_ID:Year | 5.224E-07 | 3.491E-08 | 14.963 | 36.45 |  |
|  | units!R | 2.084E-07 | 8.920E-09 | 23.365 | 14.54 |  |
| slp<sub>SER~T</sub> | Gen_ID | 7.348E-08 | 7.184E-09 | 10.229 | 55.00 | 0.81 |
|  | Gen_ID:Year | 4.516E-08 | 2.925E-09 | 15.440 | 33.80 |  |
|  | units!R | 1.495E-08 | 6.396E-10 | 23.372 | 11.19 |  |
| FH | Gen_ID | 1.226E-02 | 9.798E-04 | 12.511 | 92.24 | 0.98 |
|  | Gen_ID:Year | 5.890E-04 | 4.568E-05 | 12.893 | 4.43 |  |
|  | units!R | 4.417E-04 | 1.890E-05 | 23.371 | 3.32 |  |
| GDD<sub>15</sub> | Gen_ID | 1.226E+03 | 1.171E+02 | 10.471 | 56.64 | 0.82 |
|  | Gen_ID:Year | 6.241E+02 | 4.370E+01 | 14.283 | 28.84 |  |
|  | units!R | 3.144E+02 | 1.345E+01 | 23.365 | 14.53 |  |
| GDD<sub>95</sub> | Gen_ID | 1.190E+03 | 1.120E+02 | 10.624 | 56.84 | 0.84 |
|  | Gen_ID:Year | 4.953E+02 | 3.958E+01 | 12.515 | 23.66 |  |
|  | units!R | 4.081E+02 | 1.747E+01 | 23.365 | 19.50 |  |
| time<sub>SE</sub> | Gen_ID | 5.844E+00 | 8.095E-01 | 7.219 | 27.84 | 0.59 |
|  | Gen_ID:Year | 9.481E+00 | 6.911E-01 | 13.717 | 45.16 |  |
|  | units!R | 5.668E+00 | 2.425E-01 | 23.370 | 27.00 |  |
| GDD<sub>SE</sub> | Gen_ID | 5.665E+02 | 8.544E+01 | 6.631 | 24.14 | 0.54 |
|  | Gen_ID:Year | 1.067E+03 | 8.007E+01 | 13.326 | 45.46 |  |
|  | units!R | 7.134E+02 | 3.052E+01 | 23.375 | 30.40 |  |
| heading<sub>GDD</sub> | Gen_ID | 1.742E+03 | 1.481E+02 | 11.764 | 80.24 | 0.92 |
|  | units!R | 4.290E+02 | 2.380E+01 | 18.028 | 19.76 |  |

**Table 2:** Type-II analysis of variance table for the linear model^1^ used to predict final canopy height based on temperature response (slp_SER^~^T_) vigour (int_SER^~^T_) and stem elongation duration (GDD_SE_). *Rht-B1, Rht-D1* and *Ppd-D1* alleles and all two-way interactions^2^ were included to test for possible confounding of temperature response with final height or photoperiod. The model was applied on the 3-Year BLUES of all genotypes with available allelic data of the respective genes (n = 301)

| predictor | Sum Sq | Df | F value | Pr(>F) |  |
| --- | --- | --- | --- | --- | --- |
| $slp_{SER \sim T}$ | 7.63E-01 | 1 | 2.43E+03 | 5.08E-140 | *** |
| $int_{SER \sim T}$ | 5.91E-01 | 1 | 1.88E+03 | 2.57E-126 | *** |
| $GDD_{SE}$ | 1.29E-01 | 1 | 4.12E+02 | 5.74E-57 | *** |
| *Rht-B1* | 3.16E-03 | 1 | 1.01E+01 | 1.69E-03 | ** |
| *Rht-D1* | 3.53E-03 | 1 | 1.12E+01 | 9.13E-04 | *** |
| *Ppd-D1* | 1.48E-06 | 1 | 4.71E-03 | 9.45E-01 |  |
| $slp_{SER \sim T} : int_{SER \sim T}$ | 2.56E-04 | 1 | 8.16E-01 | 3.67E-01 | |
| $slp_{SER \sim T} : GDD_{SE}$ | 2.31E-05 | 1 | 7.36E-02 | 7.86E-01 | |
| $int_{SER \sim T} : GDD_{SE}$ | 9.76E-06 | 1 | 3.11E-02 | 8.60E-01 | |
| *Rht-B1*:*Ppd-D1* | 8.94E-04 | 1 | 2.85E+00 | 9.26E-02 | . |
| *Rht-D1*:*Ppd-D1* | 1.42E-03 | 1 | 4.51E+00 | 3.45E-02 | * |
| $slp_{SER \sim T} : Rht-B1$ | 4.01E-06 | 1 | 1.28E-02 | 9.10E-01 | |
| $slp_{SER \sim T} : Rht-D1$ | 1.43E-06 | 1 | 4.56E-03 | 9.46E-01 | |
| $slp_{SER \sim T} : Ppd-D1$ | 5.53E-04 | 1 | 1.76E+00 | 1.85E-01 | |
| $int_{SER \sim T} : Rht-B1$ | 9.01E-04 | 1 | 2.87E+00 | 9.13E-02 | . |
| $int_{SER \sim T} : Rht-D1$ | 7.02E-04 | 1 | 2.23E+00 | 1.36E-01 | |
| $int_{SER \sim T} : Ppd-D1$ | 7.70E-04 | 1 | 2.45E+00 | 1.19E-01 | |
| $GDD_{SE} : Rht-B1$ | 1.71E-05 | 1 | 5.46E-02 | 8.15E-01 | |
| $GDD_{SE} : Rht-D1$ | 7.56E-04 | 1 | 2.41E+00 | 1.22E-01 | |
| $GDD_{SE} : Ppd-D1$ | 4.48E-05 | 1 | 1.43E-01 | 7.06E-01 | |
| Residuals | 8.79E-02 | 280 |  |  |  |

<sup>1</sup>see eq. 6
<sup>2</sup>The interaction effect of *Rht-B1* and *Rht-D1* was dropped due to singularity.

**Fig. 2:**
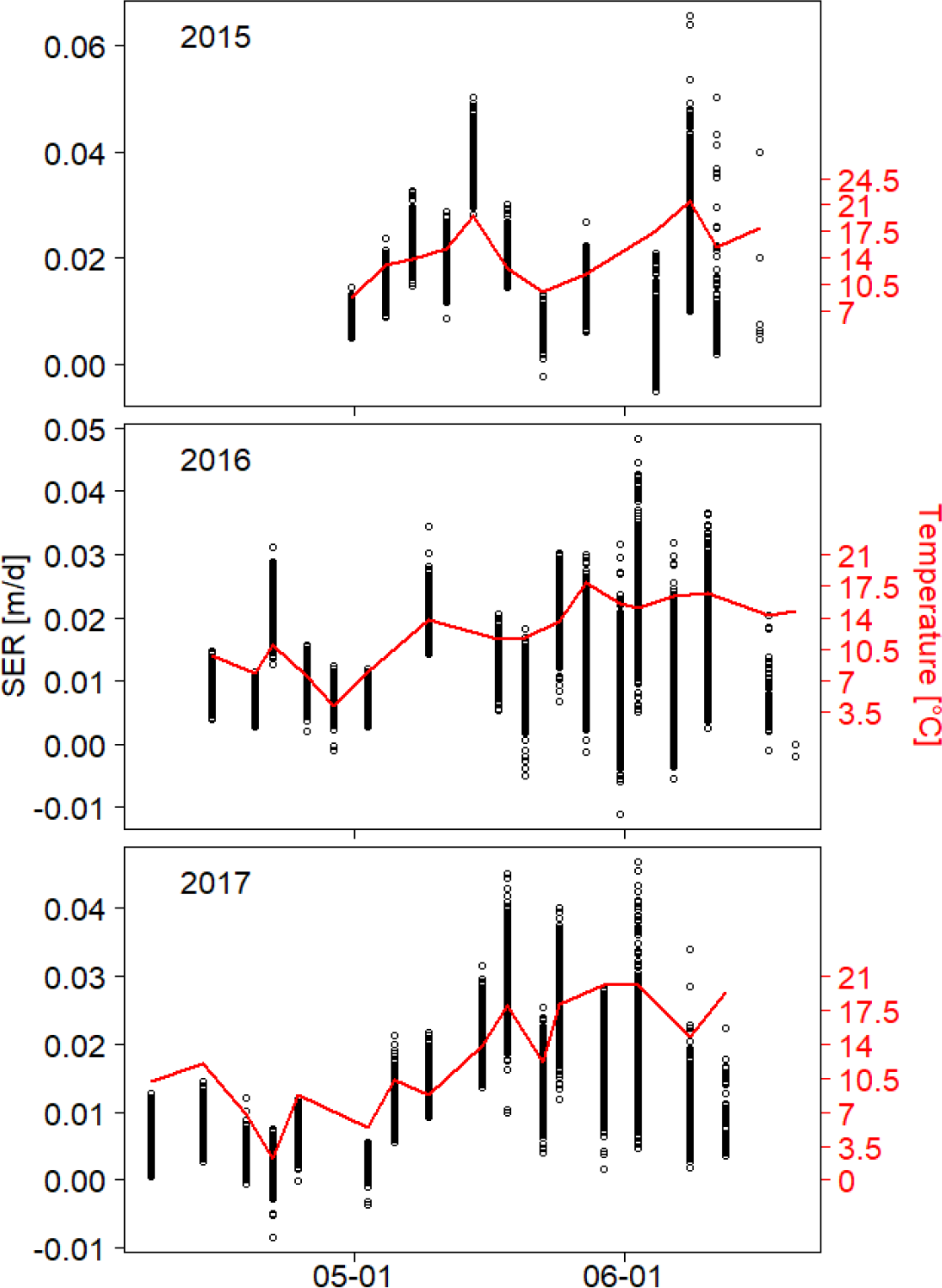
Relationship between stem elongation rate (SER) and temperature. Plot based SER raw data (n > 700/a) of > 330 genotypes (black dots) as well as temperature (solid red line) is plotted against calendar time for the years 2015-2017

**Fig. 3:**
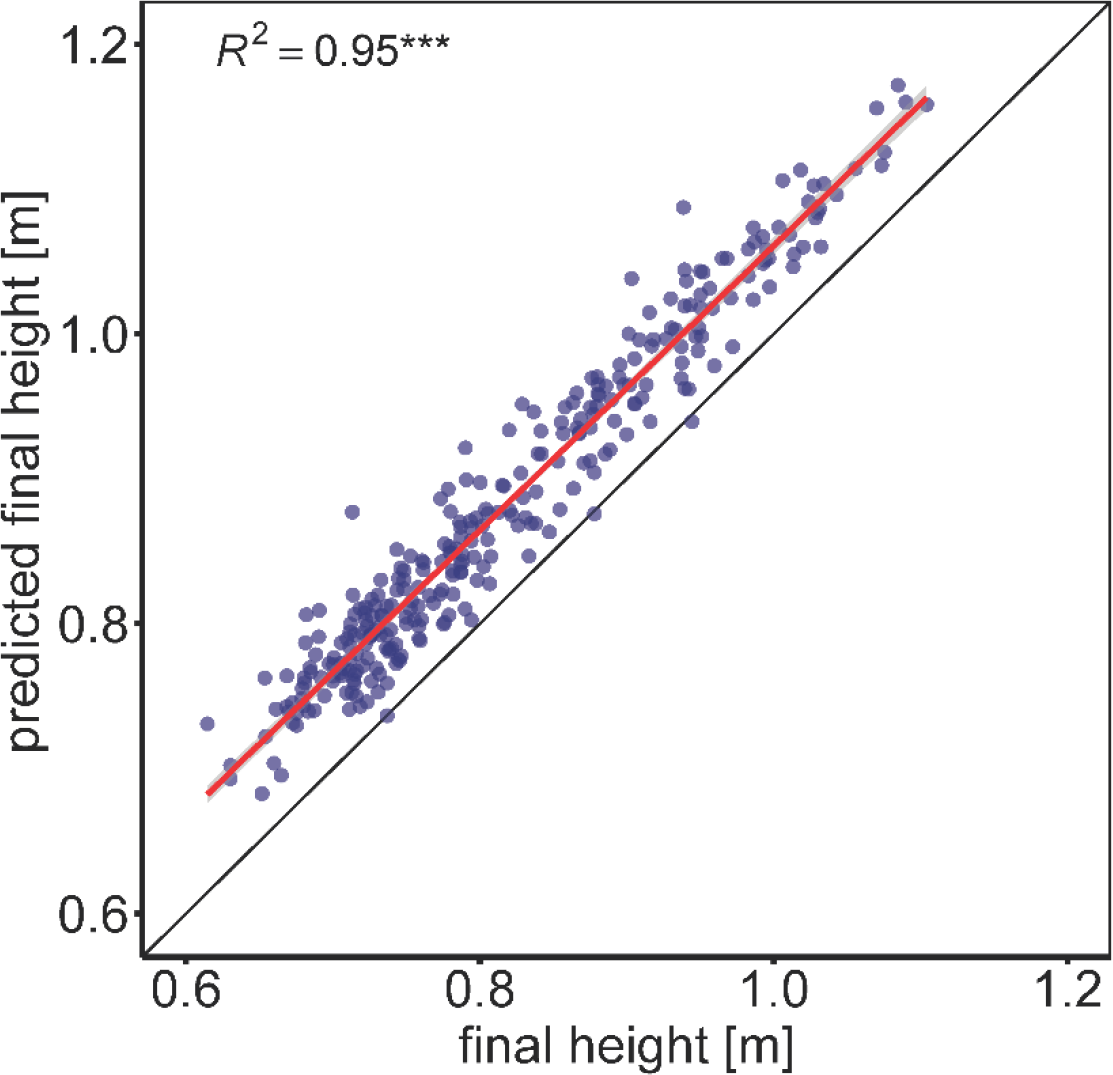
Prediction of final height based on temperature response (slp_SER^~^T_) vigour (int_SER^~^T_) and stem elongation duration (GDD_SE_). *Rht-B1, Rht-D1* and *Ppd-D1* alleles and all two-way interactions were included to test for possible confounding of temperature response with final height or photoperiod (see eq. 6 and Table 2). The model was applied on the 3-Year BLUES of all genotypes with available allelic data of the respective genes (n = 301).

### Phenology, temperature response and final height were positively correlated

To evaluate the relationships between the traits measured, Pearson correlation coefficients were calculated for each trait pair. If not indicated otherwise, the reported correlations were highly significant (*P* < 0.001)

Positive correlations were found among GDD_15_, GDD_95_ and FH (0.36 <= *r* <= 0.64, Fig. 4), indicating that taller genotypes were generally later in their development towards FH. Temperature response (slp_SER^~^T_) and vigour (int_SER^~^T_) also showed a strong, positive relationship with FH (*r* = 0.85 and *r* = 0.65, respectively). However, only temperature response correlated with GDD_15_ and GDD_95_ (*r* = 0.63 and *r* = 59, respectively), whereas vigour did not (*r* < 0.26, Fig. S3).

**Fig. 4.**
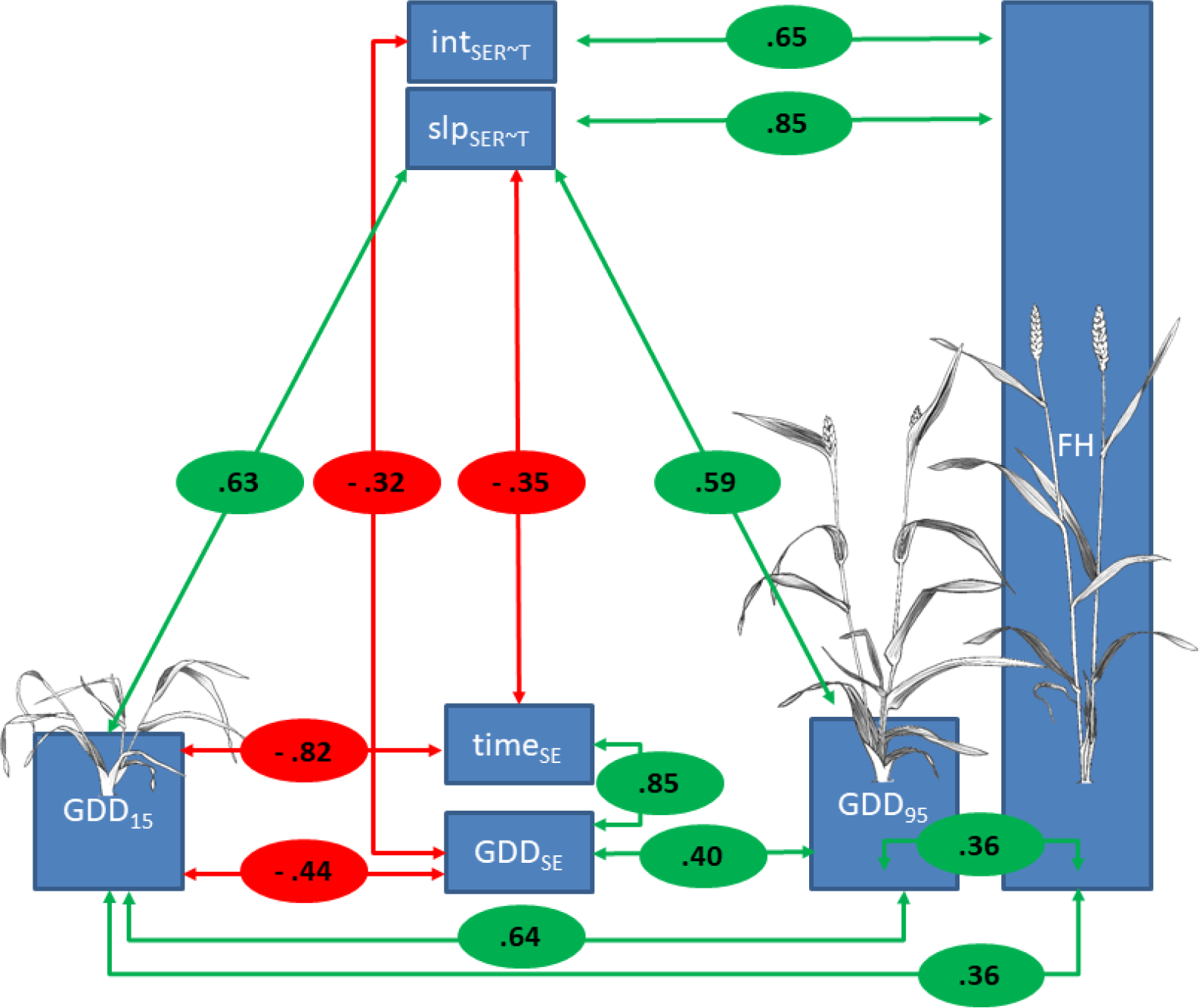
Key correlations among investigated traits. Pearson correlation coefficients between respective traits are given in red and green circles, where red denotes a negative correlation and green denotes a positive correlation. Weak correlations (*r* < 0.3) are shown in the complete correlation matrix Fig. S3. Illustrations of GDD_15_, GDD_95_ and FH were taken from Schürch *et al*. (2018).

As expected, SE duration in thermal time (GDD_SE_) was negatively correlated with GDD_15_ (*r* = −0.44) and positively correlated with GDD_95_ (*r* = 0.4). But, GDD_SE_ did not correlate with FH (*r* = – 0.01, *P* = 0.874) or temperature response (*r* = 0.006, *P* = 0.285), although GDD_SE_ negatively correlated with vigour (*r* = −0.32). In contrast, SE duration in calendar days (time_SE_) was negatively correlated with temperature response (*r* = −0.35) and GDD_15_ (*r* = −0.82), indicating a longer SE phase for earlier genotypes. Heading showed strong positive correlations with GDD 15 (*r* = 0.61) and GDD_95_ (*r* = 0.71) and a weak correlation to temperature response (*r* = 0.29). Furthermore, heading correlated negatively with int_SER^~^T_ (*r* = −0.41) and showed no correlation to FH (*r* = 0, *P* = 0.934). Other weak correlations (*r* < 0.3), that are not discussed, are shown in Fig. S3.

### Linkage disequilibrium and population structure

Prior to MTA analysis we evaluated population structure and LD. Principal component analysis of the marker genotypes revealed no distinct substructure in the investigated population. The biplot of the first two principal components showed no apparent clusters, with the first component explaining 8% and the second component explaining 3.3% of the variation in the population (Fig. S4). This is consistent with prior work using the same population (Kollers *et al*., 2013; Yates *et al*., 2019). On average across all chromosomes, LD decayed below an *r^2^* of 0.2 at a distance of 9 MB. There was however considerable variation in this threshold among the single chromosomes (Table S2).

### Association study

Genome-wide association results differed markedly depending on the applied model. Using a MLM with kinship matrix and PCA as covariates resulted in no significant MTA for any trait (Fig. S5). In contrast, the GLM using the haplotype method on temperature response yielded 2958 significant MTA for α < 0.05 and 1852 MTA for α < 0.001 respectively (Fig. S6). However, investigation of the respective QQ-plots showed large *P*-value inflation in the haplotype method whereas the *P*-values were slightly deflated when using the MLM approach (Fig. S5, Fig. S6). In contrast, with FamCPU the QQ-plots (Fig 5) showed no *P*-value inflation, except for some markers. This pattern is expected, if population structure is appropriately controlled. Therefore, FarmCPU was chosen to be the most appropriate method for the given data, despite identifying less significant MTA.

**Fig. 5.**
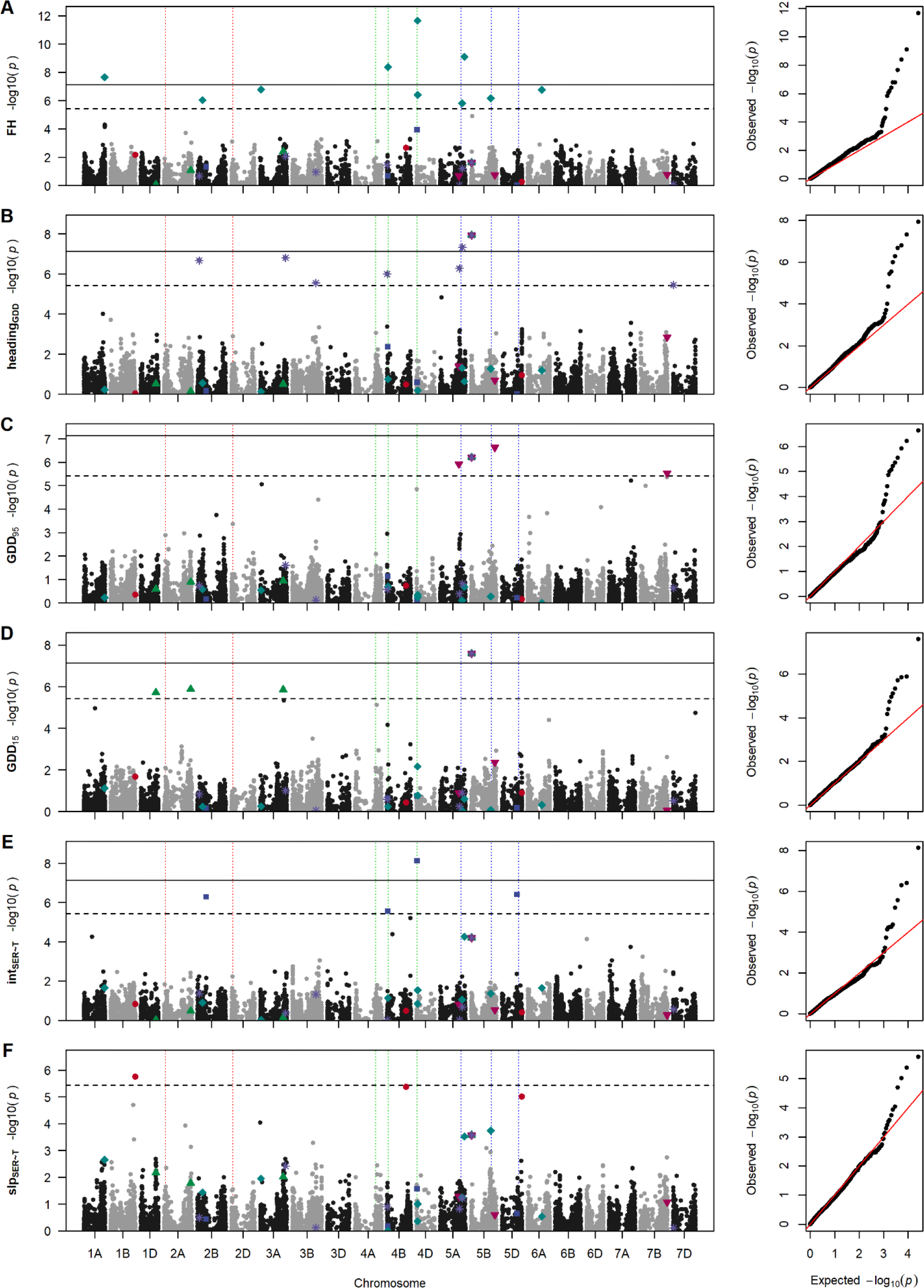
Manhattan plots and quantile-quantile plots depicting the GWAS results using FarmCPU for final height (FH, A), growing degree days until heading (heading_GDD_, B); end (GDD_95_, C) and start (GDD_15_, D) of stem elongation; vigour-related intercept (int_SER^~^T_, E) and temperature-related slope (slp_SER^~^T_, F) of stem elongation in response to temperature. Horizontal lines mark the Bonferroni corrected significance threshold for *P* < 0.05 (dashed line) and *P* < 0.001 (solid line). Vertical dotted lines mark the positions of *Ppd-1* on chromosomes 2A and 2D (red), *Rht-1* on chromosomes 4A-4D (green) and *Vrn-1* on chromosomes 5A-5D. Significant marker trait associations for slp_SER^~^T_ (red dots), int_SER^~^T_ (blue squares), GDD_15_ (green up-facing triangles), GDD_95_ (magenta down-facing triangles), heading (violet asterisks) and FH (turquois diamonds) are highlighted in all manhattan plots.

As a positive control we compared our FH data and associated markers with data made by Zanke et al (2014b) who used the same population and SNP chip in field experiments in France and Germany. Final canopy height correlated strongly between the two studies (*r* = 0.95), which is in accordance with the high heritability of the trait. In this study, we found ten significant MTA for FH (Table 3, Fig. 5). Zanke et al. (2014b) reported 280 significant MTA for FH across several environments. Of these, only marker RAC875_rep_c105718_585 on chromosome 4D overlapped with the MTA found in this study. However, by considering flanking markers, we found that of the remaining nine significant MTA for FH, four were in LD with MTA found by Zanke et al. (2014b; Table S3). The significant MTA found for FH in this study are near known genes controlling FH. For example, Tdurum_contig64772_417, is 4 MB upstream of *Rht*-B1 and RAC875_rep_c105718_585, is 7 MB downstream of *Rht*-D1 on their respective group 4 chromosomes.

**Table 3:** Marker-trait associations for slp_SER^~^T_, int_SER^~^T_, GDD_15_, GDD_95_ heading_GDD_ and FH, including *P*-value, minor allele frequency and allelic effect estimate.

| Trait | SNP | Chr | Position | $P$-value | maf | effect |
| --- | --- | --- | --- | --- | --- | --- |
| $slp_{SER\sim T}$ | wsnp_Ex_c1597_3045682 | 1B | 688'283'256 | 1.76E-06 | 0.19 | -6.05E-05 |
|  | CAP7_c10839_300 | 4B | 533'724'424 | 4.17E-06 | 0.24 | -5.07E-05 |
|  | IAAV7104 | 5D | 553'678'522 | 9.67E-06 | 0.13 | -6.02E-05 |
| $int_{SER\sim T}$ | RAC875_s109189_188 | 2B | 248'149'774 | 5.08E-07 | 0.42 | 1.73E-04 |
|  | Ku_c63300_1309 | 4B | 21'556'672 | 2.73E-06 | 0.10 | -2.99E-04 |
|  | Kukri_rep_c68594_530 | 4D | 12'773'259 | 7.47E-09 | 0.40 | -2.30E-04 |
|  | Kukri_c6477_696 | 5D | 423'502'809 | 3.89E-07 | 0.21 | -2.03E-04 |
| $GDD_{15}$ | wsnp_Ex_c12447_19847242 | 1D | 416'456'386 | 1.91E-06 | 0.46 | 6.89E+00 |
|  | Tdurum_contig47508_250 | 2A | 754'339'235 | 1.31E-06 | 0.21 | 9.41E+00 |
|  | Kukri_c55381_67 | 3A | 648'868'234 | 1.38E-06 | 0.17 | -1.00E+01 |
|  | Excalibur_c74858_243 | 5B | 13'190'663 | 2.50E-08 | 0.47 | -7.88E+00 |
| $GDD_{95}$ | Excalibur_c49597_579 | 5A | 521'934'666 | 1.19E-06 | 0.42 | -6.58E+00 |
|  | Excalibur_c74858_243 | 5B | 13'190'663 | 6.15E-07 | 0.47 | -6.14E+00 |
|  | Tdurum_contig44115_561 | 5B | 669'897'388 | 2.31E-07 | 0.13 | -1.02E+01 |
|  | RAC875_c38693_319 | 7B | 740'056'880 | 2.87E-06 | 0.20 | 7.51E+00 |
| $heading_{GDD}$ | RAC875_c12766_461 | 2B | 47'430'682 | 2.10E-07 | 0.39 | -7.77E+00 |
|  | Kukri_rep_c106620_208 | 3A | 714'300'397 | 1.55E-07 | 0.08 | 1.47E+01 |
|  | BS00022611_51 | 3B | 659'787'924 | 2.83E-06 | 0.12 | 8.75E+00 |
|  | IAAV7221 | 4B | 2'036'611 | 9.96E-07 | 0.07 | -1.29E+01 |
|  | BS00000365_51 | 5A | 538'000'573 | 5.19E-07 | 0.44 | -7.65E+00 |
|  | IACX2540 | 5A | 619'684'943 | 4.73E-08 | 0.35 | 9.39E+00 |
|  | Excalibur_c74858_243 | 5B | 13'190'663 | 1.16E-08 | 0.47 | -8.67E+00 |
|  | Excalibur_c46904_84 | 7D | 5'198'912 | 3.55E-06 | 0.13 | -9.46E+00 |
| $FH$ | Excalibur_c85499_232 | 1A | 582'219'427 | 2.23E-08 | 0.11 | 2.52E-02 |
|  | wsnp_Ku_c11665_18999583 | 2B | 139'070'721 | 9.07E-07 | 0.13 | 2.08E-02 |
|  | Kukri_c49280_230 | 3A | 20'134'735 | 1.63E-07 | 0.08 | 2.90E-02 |
|  | Tdurum_contig64772_417 | 4B | 26'491'482 | 4.18E-09 | 0.07 | 3.56E-02 |
|  | RAC875_rep_c105718_585 | 4D | 25'989'162 | 2.20E-12 | 0.38 | -2.56E-02 |
|  | BS00036421_51 | 4D | 32'347'318 | 3.96E-07 | 0.37 | -1.58E-02 |
|  | RAC875_c8231_1578 | 5A | 613'588'253 | 1.52E-06 | 0.43 | 1.44E-02 |
|  | wsnp_Ku_rep_c71232_70948744 | 5A | 679'663'586 | 7.93E-10 | 0.47 | -2.21E-02 |
|  | BS00109560_51 | 5B | 556'182'591 | 6.91E-07 | 0.46 | -1.56E-02 |
|  | BS00022120_51 | 6A | 396'301'470 | 1.68E-07 | 0.24 | -2.01E-02 |

### Temperature-response loci are independent of vigour loci

For slp_SER^~^T_ we detected one significant (LOD = 5.75) MTA on chromosome 1B (wsnp_Ex_c1597_3045682) and two almost significant (LOD = 5.38 / LOD = 5.01) MTA on chromosomes 4B (CAP7_c10839_300) and 5D (IAAV7104), respectively (Fig. 5). All associated markers for slp_SER^~^T_ yielded small but significant allelic effects ranging from −0.061 mm °C^−1^d^−1^ to −0.051 mm °C^−1^d^−1^ (Table 3). The GWAS for int_SER^~^T_ yielded four significant MTA on chromosomes 2B, 4B, 4D and 5D respectively (Table 3, Fig. 5). Start and end of SE yielded four MTA each and heading yielded eight MTA (Table 3, Fig. 5).

Comparing the GWAS results for temperature response, vigour, FH, GDD_15_, GDD_95_ and heading revealed no common quantitative trait loci (QTL) between slp_SER^~^T_ and any other trait. Only one marker (Excalibur_c74858_243) was significantly associated with both GDD_15_ and GDD_95_, as well as heading. The lack of overlap, of MTA, between temperature response, vigour and timing of critical stages indicate they are genetically independent. However, there is a genetic connection between vigour and FH on the one hand and between the start and end of SE and heading on the other.

To identify potential causative genes underlying the QTL, we searched the reference genome annotation around the respective QTL intervals. For temperature response we found an increased presence of genes or gene homologues involved in the flowering pathway, i.e. *EARLY FLOWERING 3, FRIGIDA* and *CONSTANS* (Table 4). Around the QTL associated with vigour the annotation showed genes associated with growth (i.e. *GRAS, CLAVATA, BSU1, ARGONAUTE*) as well as developmental progress (i.e. *Tesmin/TSO1-like CXC domain, BEL1, AGAMOUS* (Table 5). Importantly, we found *GAI-like protein 1* 6MB upstream of marker Kukri_rep_c68594_530, which we identified as *Rht-D1* by blasting the *Rht*-D1 sequence (GeneBank ID AJ242531.1) against the annotated reference genome. Genes putatively underlying the QTL for heading, GDD_15_ and GDD_95_ are listed in the supplementary tables S4 – S6. As expected, genes or gene homologues associated with the flowering pathway were found in the vicinity of the MTA for heading. The common QTL for heading, GDD_15_ and GDD_95_ on chromosome 5B (Excalibur_c74858_243) was found to be 6.6 MB upstream of *FLOWERING LOCUS T* (Tables S4-S6). Other flowering associated genes found near the heading QTL were CONSTANS, *FRIGIDA*, and a *FLOWERING LOCUS C* associated protein (Table S4). Moreover, a number of putative response regulators as well as genes putatively involved in light control of development (i.e. FAR1-RELATED SEQUENCE; Lin and Wang, 2004) were found near the heading QTL (Table S4). The remaining QTL for GDD_95_ were near genes associated with developmental progress and flowering, e.g. *AGAMOUS, MEI2-like 1, HAPPLESS 2* and *BEL1* (Table S5). Genes near the remaining QTL for GDD_15_ were associated with developmental progress (*i.e. FLOWERING LOCUS T, BEL1, TERMINAL EAR1-like*, and *FAR1-RELATED SEQUENCE*) as well as growth (*i.e. CLAVATA* and DELLA; Table S6).

**Table 4:** Selected putative candidate genes for temperature response (slp_SER^~^T_) from the IWGSC reference genome annotation.

| Chr | SNP [Position] | r.start | r.end | Gene | description | distance |
| --- | --- | --- | --- | --- | --- | --- |
| chr1B | wsnp_Ex_c1597_3045682<br>[688'283'256] | 688'282'509 | 688'286'431 | TraesCS1B01G480600 | winged-helix DNA-binding transcription factor family protein | 747 |
|  |  | 688'352'414 | 688'354'696 | TraesCS1B01G480700 | HMG-Y-related protein A | -69'158 |
|  |  | 687'710'716 | 687'719'885 | TraesCS1B01G480100 | Argonaute | 572'540 |
|  |  | 687'128'952 | 687'135'442 | TraesCS1B01G479200 | Zinc finger protein CONSTANS | 1'154'304 |
|  |  | 687'078'233 | 687'084'562 | TraesCS1B01G479000 | Zinc finger protein CONSTANS | 1'205'023 |
|  |  | 686'928'468 | 686'931'886 | TraesCS1B01G478700 | Zinc finger protein CONSTANS | 1'354'788 |
|  |  | 686'749'516 | 686'755'405 | TraesCS1B01G478100 | WD-repeat protein, putative | 1'533'740 |
|  |  | 685'645'287 | 685'649'392 | TraesCS1B01G477400 | Early flowering 3 | 2'637'969 |
| chr4B | CAP7_c10839_300<br>[533'724'424] | 537'474'959 | 537'479'867 | TraesCS4B01G266000 | Protein FRIGIDA | -3'750'535 |
|  |  | 541'363'317 | 541'365'139 | TraesCS4B01G267700 | Protein upstream of flc | -7'638'893 |
|  |  | 542'582'729 | 542'583'265 | TraesCS4B01G268300 | MADS transcription factor | -8'858'305 |
| chr5D | IAAV7104<br>[553'678'522] | 554'357'761 | 554'360'305 | TraesCS5D01G544800 | FRIGIDA-like protein, putative | -679'239 |
|  |  | 554'467'487 | 554'472'596 | TraesCS5D01G545100 | Transducin/WD-like repeat-protein | -788'965 |
|  |  | 556'226'523 | 556'234'480 | TraesCS5D01G548800 | Transducin/WD-like repeat-protein | -2'548'001 |

**Table 5:**
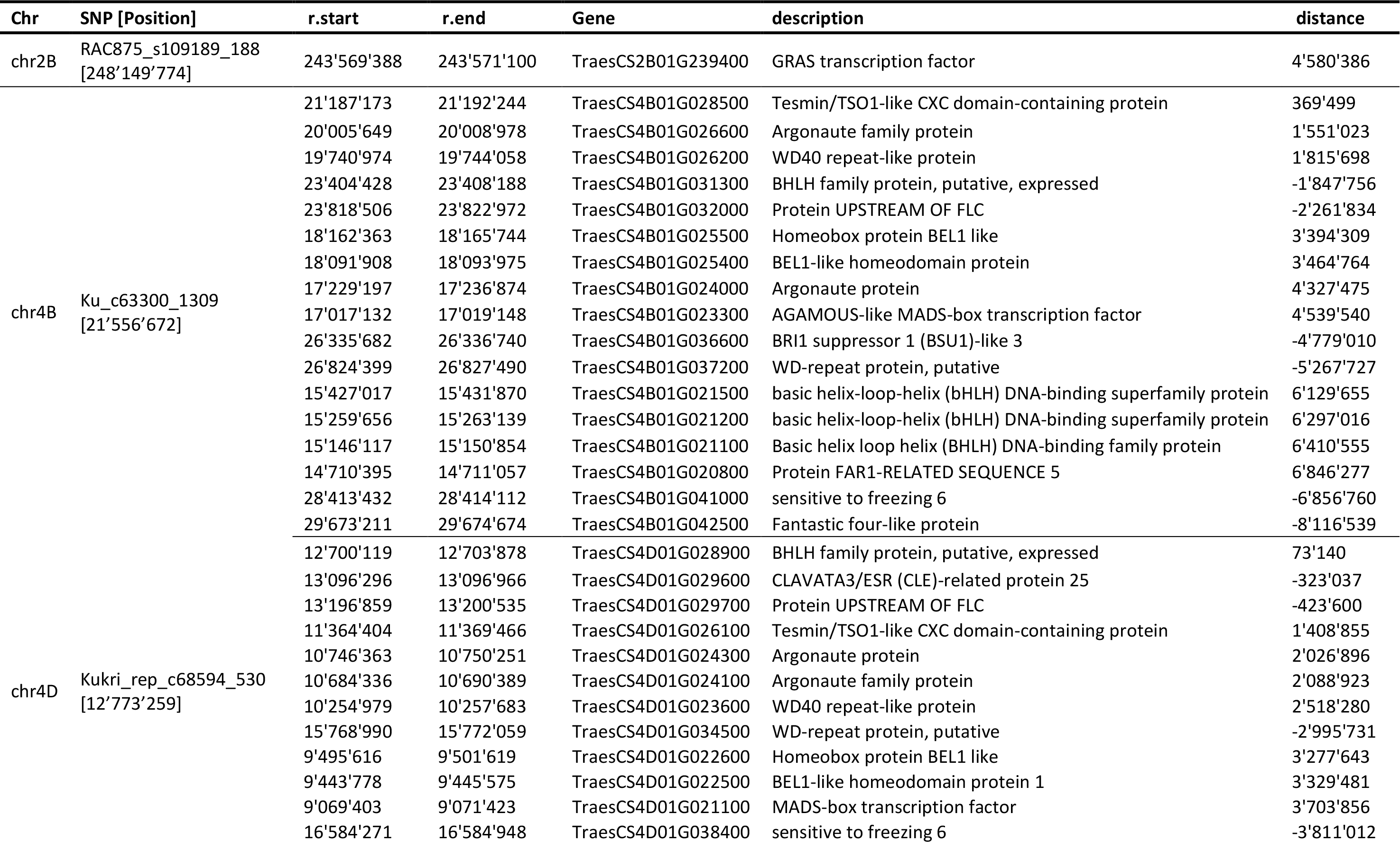
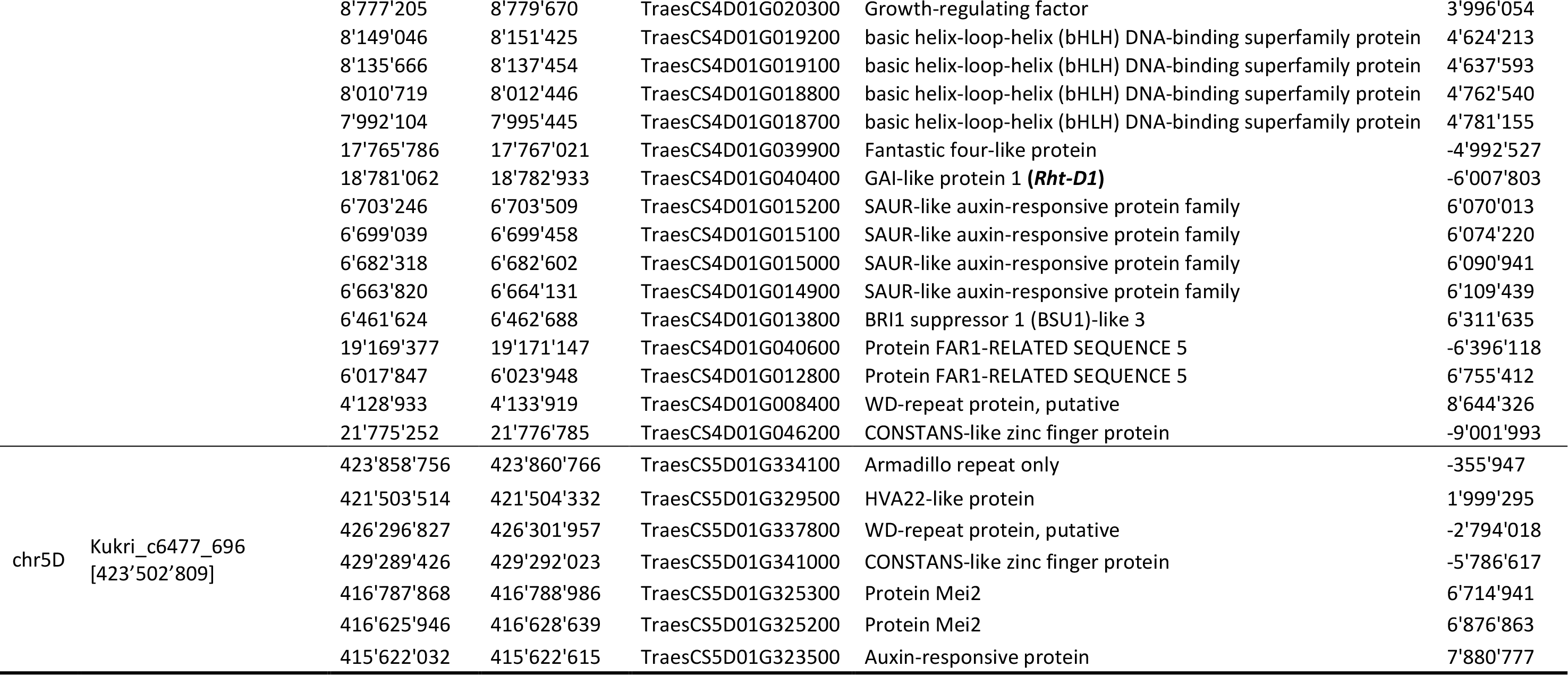
Selected putative candiate genes for vigour (int_SER^~^T_) of temperature response from the IWGSC reference genome annotation.

| Chr | SNP [Position] | r.start | r.end | Gene | description | distance |
| --- | --- | --- | --- | --- | --- | --- |
| chr2B | RAC875_s109189_188<br>[248'149'774] | 243'569'388 | 243'571'100 | TraesCS2B01G239400 | GRAS transcription factor | 4'580'386 |
| chr4B | Ku_c63300_1309<br>[21'556'672] | 21'187'173 | 21'192'244 | TraesCS4B01G028500 | Tesmin/TSO1-like CXC domain-containing protein | 369'499 |
|  |  | 20'005'649 | 20'008'978 | TraesCS4B01G026600 | Argonaute family protein | 1'551'023 |
|  |  | 19'740'974 | 19'744'058 | TraesCS4B01G026200 | WD40 repeat-like protein | 1'815'698 |
|  |  | 23'404'428 | 23'408'188 | TraesCS4B01G031300 | BHLH family protein, putative, expressed | -1'847'756 |
|  |  | 23'818'506 | 23'822'972 | TraesCS4B01G032000 | Protein UPSTREAM OF FLC | -2'261'834 |
|  |  | 18'162'363 | 18'165'744 | TraesCS4B01G025500 | Homeobox protein BEL1 like | 3'394'309 |
|  |  | 18'091'908 | 18'093'975 | TraesCS4B01G025400 | BEL1-like homeodomain protein | 3'464'764 |
|  |  | 17'229'197 | 17'236'874 | TraesCS4B01G024000 | Argonaute protein | 4'327'475 |
|  |  | 17'017'132 | 17'019'148 | TraesCS4B01G023300 | AGAMOUS-like MADS-box transcription factor | 4'539'540 |
|  |  | 26'335'682 | 26'336'740 | TraesCS4B01G036600 | BRI1 suppressor 1 (BSU1)-like 3 | -4'779'010 |
|  |  | 26'824'399 | 26'827'490 | TraesCS4B01G037200 | WD-repeat protein, putative | -5'267'727 |
|  |  | 15'427'017 | 15'431'870 | TraesCS4B01G021500 | basic helix-loop-helix (bHLH) DNA-binding superfamily protein | 6'129'655 |
|  |  | 15'259'656 | 15'263'139 | TraesCS4B01G021200 | basic helix-loop-helix (bHLH) DNA-binding superfamily protein | 6'297'016 |
|  |  | 15'146'117 | 15'150'854 | TraesCS4B01G021100 | Basic helix loop helix (BHLH) DNA-binding family protein | 6'410'555 |
|  |  | 14'710'395 | 14'711'057 | TraesCS4B01G020800 | Protein FAR1-RELATED SEQUENCE 5 | 6'846'277 |
|  |  | 28'413'432 | 28'414'112 | TraesCS4B01G041000 | sensitive to freezing 6 | -6'856'760 |
|  |  | 29'673'211 | 29'674'674 | TraesCS4B01G042500 | Fantastic four-like protein | -8'116'539 |
| chr4D | Kukri_rep_c68594_530<br>[12'773'259] | 12'700'119 | 12'703'878 | TraesCS4D01G028900 | BHLH family protein, putative, expressed | 73'140 |
|  |  | 13'096'296 | 13'096'966 | TraesCS4D01G029600 | CLAVATA3/ESR (CLE)-related protein 25 | -323'037 |
|  |  | 13'196'859 | 13'200'535 | TraesCS4D01G029700 | Protein UPSTREAM OF FLC | -423'600 |
|  |  | 11'364'404 | 11'369'466 | TraesCS4D01G026100 | Tesmin/TSO1-like CXC domain-containing protein | 1'408'855 |
|  |  | 10'746'363 | 10'750'251 | TraesCS4D01G024300 | Argonaute protein | 2'026'896 |
|  |  | 10'684'336 | 10'690'389 | TraesCS4D01G024100 | Argonaute family protein | 2'088'923 |
|  |  | 10'254'979 | 10'257'683 | TraesCS4D01G023600 | WD40 repeat-like protein | 2'518'280 |
|  |  | 15'768'990 | 15'772'059 | TraesCS4D01G034500 | WD-repeat protein, putative | -2'995'731 |
|  |  | 9'495'616 | 9'501'619 | TraesCS4D01G022600 | Homeobox protein BEL1 like | 3'277'643 |
|  |  | 9'443'778 | 9'445'575 | TraesCS4D01G022500 | BEL1-like homeodomain protein 1 | 3'329'481 |
|  |  | 9'069'403 | 9'071'423 | TraesCS4D01G021100 | MADS-box transcription factor | 3'703'856 |
|  |  | 16'584'271 | 16'584'948 | TraesCS4D01G038400 | sensitive to freezing 6 | -3'811'012 |
|  |  | 8'777'205 | 8'779'670 | TraesCS4D01G020300 | Growth-regulating factor | 3'996'054 |
|  |  | 8'149'046 | 8'151'425 | TraesCS4D01G019200 | basic helix-loop-helix (bHLH) DNA-binding superfamily protein | 4'624'213 |
|  |  | 8'135'666 | 8'137'454 | TraesCS4D01G019100 | basic helix-loop-helix (bHLH) DNA-binding superfamily protein | 4'637'593 |
|  |  | 8'010'719 | 8'012'446 | TraesCS4D01G018800 | basic helix-loop-helix (bHLH) DNA-binding superfamily protein | 4'762'540 |
|  |  | 7'992'104 | 7'995'445 | TraesCS4D01G018700 | basic helix-loop-helix (bHLH) DNA-binding superfamily protein | 4'781'155 |
|  |  | 17'765'786 | 17'767'021 | TraesCS4D01G039900 | Fantastic four-like protein | -4'992'527 |
|  |  | 18'781'062 | 18'782'933 | TraesCS4D01G040400 | GAI-like protein 1 (**Rht-D1**) | -6'007'803 |
|  |  | 6'703'246 | 6'703'509 | TraesCS4D01G015200 | SAUR-like auxin-responsive protein family | 6'070'013 |
|  |  | 6'699'039 | 6'699'458 | TraesCS4D01G015100 | SAUR-like auxin-responsive protein family | 6'074'220 |
|  |  | 6'682'318 | 6'682'602 | TraesCS4D01G015000 | SAUR-like auxin-responsive protein family | 6'090'941 |
|  |  | 6'663'820 | 6'664'131 | TraesCS4D01G014900 | SAUR-like auxin-responsive protein family | 6'109'439 |
|  |  | 6'461'624 | 6'462'688 | TraesCS4D01G013800 | BRI1 suppressor 1 (BSU1)-like 3 | 6'311'635 |
|  |  | 19'169'377 | 19'171'147 | TraesCS4D01G040600 | Protein FAR1-RELATED SEQUENCE 5 | -6'396'118 |
|  |  | 6'017'847 | 6'023'948 | TraesCS4D01G012800 | Protein FAR1-RELATED SEQUENCE 5 | 6'755'412 |
|  |  | 4'128'933 | 4'133'919 | TraesCS4D01G008400 | WD-repeat protein, putative | 8'644'326 |
|  |  | 21'775'252 | 21'776'785 | TraesCS4D01G046200 | CONSTANS-like zinc finger protein | -9'001'993 |
| chr5D | Kukri_c6477_696<br>[423'502'809] | 423'858'756 | 423'860'766 | TraesCS5D01G334100 | Armadillo repeat only | -355'947 |
|  |  | 421'503'514 | 421'504'332 | TraesCS5D01G329500 | HVA22-like protein | 1'999'295 |
|  |  | 426'296'827 | 426'301'957 | TraesCS5D01G337800 | WD-repeat protein, putative | -2'794'018 |
|  |  | 429'289'426 | 429'292'023 | TraesCS5D01G341000 | CONSTANS-like zinc finger protein | -5'786'617 |
|  |  | 416'787'868 | 416'788'986 | TraesCS5D01G325300 | Protein Mei2 | 6'714'941 |
|  |  | 416'625'946 | 416'628'639 | TraesCS5D01G325200 | Protein Mei2 | 6'876'863 |
|  |  | 415'622'032 | 415'622'615 | TraesCS5D01G323500 | Auxin-responsive protein | 7'880'777 |

### Vigour, temperature response and the timing of SE affect final height

The phenotypic correlations show a strong connection between temperature response, vigour and FH as well as weaker connections between GDD_15_, GDD_95_ and FH. In order to examine this interdependency on a genetic level, we used a linear model to predict FH with the SNP alleles of the QTL for slp_SER^~^T_, int_SER^~^T_, GDD_15_ and GDD_95_ as predictors. The model was able to predict FH with an accuracy *R^2^* = 0.5, however, clusters in the data showed clear effects of *Rht-D1* and *Ppd-D1* alleles (Fig. 6A). Adding *Rht-B1, Rht-D1* and *Ppd-D1* alleles as predictors increased the prediction accuracy to *R^2^* = 0.71 (Fig 6B). There were significant contributions by QTL of all three traits, however, their effect were small compared to the obvious effects of *Rht-B1, Rht-D1 and Ppd-D1* (Table 6). Including all two-way interaction effects among the QTL, *Rht-1* and *Ppd-1* increased the prediction accuracy to *R^2^* = 0.87 (Fig. 6C) indicating a fine-tuning effect of temperature response, vigour and timing of SE towards FH.

**Table 6:** Type II analysis of variance for the prediction of final height using QTL and *Rht-B1, Rht-D1* and *Ppd-D1* alleles. SNP alleles of the QTL for temperature response (slp_SER^~^T_), vigour (int_SER^~^T_), start (GDD_15_) and end (GDD_95_) of stem elongation as well as *Rht-B1, Rht-D1* and *Ppd-D1* alleles were used as predictors in a linear model (FH = ∑ QTL slp_SER^~^T_ + ∑ QTL int_SER^~^T_ + ∑ QTL GDD_15_ + ∑ QTL GDD_95_ + *Rht-B1* + *Rht-D1* + *Ppd-D1*) without interaction effects (see Fig. 6B). The model was applied to the 3-year BLUES of all genotypes with available genotypic data (n = 300) on all predictors.

| Predictor<br>[trait.QTL.chromosome] | SNP | Sum Sq | Df | F value | Pr(>F) |  |
| --- | --- | --- | --- | --- | --- | --- |
| $slp_{SER\sim T}.1.1B$ | wsnp_Ex_c1597_3045682 | 4.20E-03 | 1 | 1.08E+00 | 2.99E-01 | |
| $slp_{SER\sim T}.2.4B$ | CAP7_c10839_300 | 7.97E-03 | 1 | 2.05E+00 | 1.53E-01 | |
| $slp_{SER\sim T}.3.5D$ | IAAV7104 | 2.81E-02 | 1 | 7.23E+00 | 7.58E-03 | ** |
| $int_{SER\sim T}.1.2B$ | RAC875_s109189_188 | 1.72E-03 | 1 | 4.42E-01 | 5.07E-01 | |
| $int_{SER\sim T}.2.4B$ | Ku_c63300_1309 | 1.64E-02 | 1 | 4.23E+00 | 4.07E-02 | * |
| $int_{SER\sim T}.3.4D$ | Kukri_rep_c68594_530 | 7.47E-03 | 1 | 1.93E+00 | 1.66E-01 | |
| $int_{SER\sim T}.4.5D$ | Kukri_c6477_696 | 6.06E-03 | 1 | 1.56E+00 | 2.13E-01 | |
| GDD<sub>15</sub>.1.1D | wsnp_Ex_c12447_19847242 | 1.41E-02 | 1 | 3.64E+00 | 5.74E-02 | . |
| GDD<sub>15</sub>.2.2A | Tdurum_contig47508_250 | 4.97E-02 | 1 | 1.28E+01 | 4.07E-04 | *** |
| GDD<sub>15</sub>.3.3A | Kukri_c55381_67 | 1.91E-02 | 1 | 4.91E+00 | 2.75E-02 | * |
| GDD<sub>15</sub>.4\|GDD<sub>95</sub>.2\|heading.7.5B | Excalibur_c74858_243 | 7.64E-08 | 1 | 1.97E-05 | 9.96E-01 |  |
| GDD<sub>95</sub>.1.5A | Excalibur_c49597_579 | 4.04E-02 | 1 | 1.04E+01 | 1.40E-03 | ** |
| GDD<sub>95</sub>.3.5B | Tdurum_contig44115_561 | 1.03E-03 | 1 | 2.65E-01 | 6.07E-01 |  |
| GDD<sub>95</sub>.4.7B | RAC875_c38693_319 | 4.28E-03 | 1 | 1.10E+00 | 2.95E-01 |  |
| *Rht-B1* |  | 3.32E-01 | 1 | 8.55E+01 | 6.02E-18 | *** |
| *Rht-D1* |  | 6.71E-01 | 1 | 1.73E+02 | 3.79E-31 | *** |
| *Ppd-D1* |  | 5.87E-02 | 1 | 1.51E+01 | 1.26E-04 | *** |
| Residuals |  | 1.09E+00 | 282 |  |  |  |

**Fig. 6:**
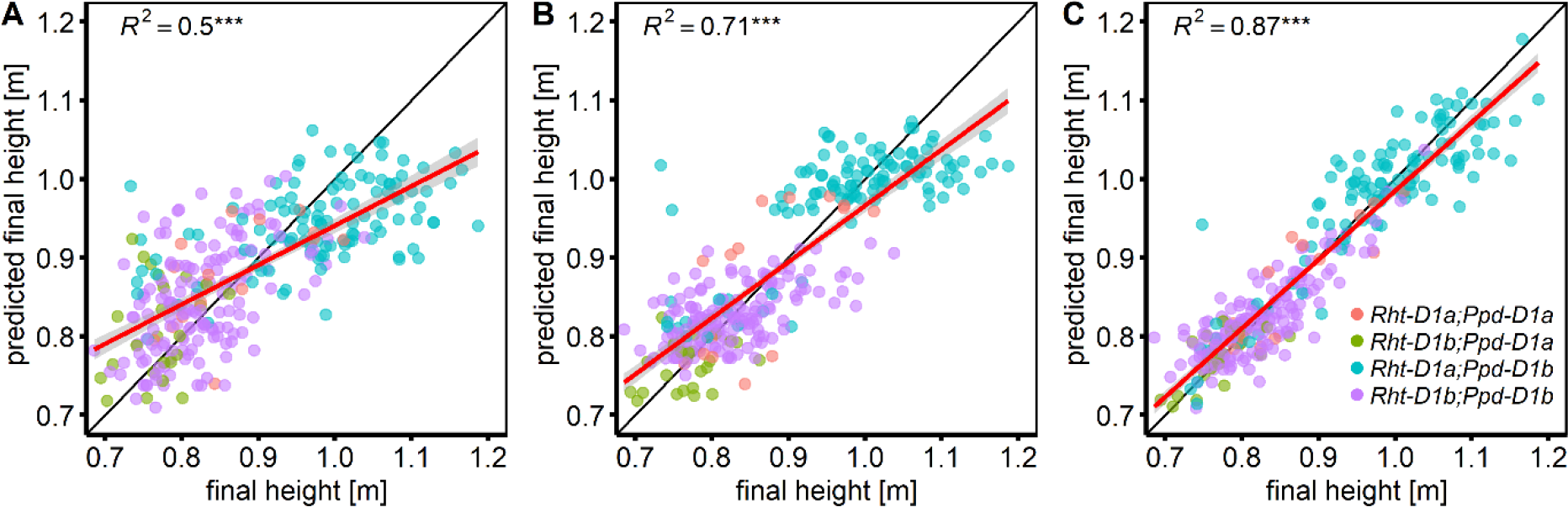
Prediction of final height based on QTL and *Rht-B1, Rht-D1* and *Ppd-D1* alleles. **A:** The SNP alleles of significantly associated QTL for temperature response (slp_SER^~^T_), vigour (int_SER^~^T_), start (GDD_15_) and end (GDD_95_) of stem elongation were used in a linear model without considering interaction effects. **B:** *Rht-B1, Rht-D1* and *Ppd-D1* alleles were added to the model used in A. **C:** All two-way interaction effects among SNP alleles and *Rht-B1, Rht-D1* and *Ppd-D1* alleles were included in the model. The models were applied to the 3-year BLUES of all genotypes with available genotypic data (n = 300) on all predictors. Colours indicate the allelic status regarding *Rht-D1* and *Ppd-D1* of the respective genotypes as depicted in panel C.

## Discussion

In this study we present a method to measure temperature response during stem elongation of wheat using high throughput phenotyping of canopy height in the field. We found a genotype-specific response of wheat to change in ambient temperature which was correlated with the timing of the developmental key stages. We decomposed this growth dynamics into a genotype-specific vigour component and temperature response component using regression models. We further related these parameters to plant height and the timing of developmental key stages.

Linear regression models were used to describe wheat growth response to temperature for leaf elongation (Nagelmüller *et al*., 2016), canopy cover (Grieder *et al*., 2015) as well as stem elongation rate (Slafer and Rawson, 1995a). Others proposed the use of a more complex, Arrhenius type of peak function to account for decreasing growth rates at supra optimal temperatures (Parent and Tardieu, 2012). However such models are mainly applicable when the temperatures experienced by the crop exceed the temperature optimum. Wheat has its temperature-optimum at around 27°C (Parent and Tardieu, 2012). Temperatures in the measured growth intervals during SE did not exceed 25°C and given the temporal resolution of the data, a simple linear model is justified (Parent *et al*., 2018).

The results of the correlation analysis show a clear connection between FH and temperature response (slp_SER^~^T_) as well as between FH and vigour (int_SER^~^T_). This is consistent with our hypothesis that FH can be described as a function of temperature-independent growth processes and as a function of temperature response during SE. Importantly, among all components, the temperature response was a significant driver of FH and also had a strong influence on the timing. Temperature response delayed the beginning of SE leading to a later start and end of the whole phase. This finding might appear counter intuitive: given the assumption that plants develop faster under higher ambient temperatures a more responsive genotype should develop faster compared to a less responsive one. Slafer and Rawson (1995b) reported an accelerated development towards floral transition with increasing temperatures up to 19°C whereas higher temperatures slowed development. In that respect, a more responsive genotype would experience a stronger delay of floral transition under warm temperatures.

In terms of their correlation to FH, the effects of the timing of start and end of SE are less distinct. FH was more a function of faster growth than a longer duration of growth, especially since genotypes with a strong temperature response had a shorter duration of SE. However, the timing of start and end of SE was linked with temperature response. Based on this result and the according correlations, it would appear that temperature response influences FH directly as well as indirectly by mediating start and end of SE. Surprisingly, we found no correlation between heading and FH despite the positive correlation of both traits with GDD_95_. A correlation between heading date and FH would be therefore expected. Previous studies reporting pleiotropic effects between plant height and heading time (Griffiths *et al*., 2010; Mo *et al*., 2018).

The question, whether these trait correlations are due to pleiotropic effects will substantially impact the breeding strategy (Chen and Lübberstedt, 2010). If the relationship between phenology, FH and temperature response would be to a large degree pleiotropic, these traits could not be independently selected. Alternative explanations are linkage and population structure. The GABI wheat panel is made of wheat varieties from different regions of Europe. As the examined traits are major drivers of adaptation to the different regions of Europe, we anticipate a very strong selection for both, temperature response as well as timing of critical stages. Even if there is no apparent population structure at neutral markers, there may be a strong population structure at selected loci with strong effect on local adaptation. Our phenotypic results showed a significant interaction effect between *Ppd-D1* and *Rht-D1* on final canopy height, indicating either a co-selection or a pleiotropic effect of *Ppd-D1*. Pleiotropic effects between height and flowering time are known for maize and rice. For example, the *DWARF8* gene of maize encoding a DELLA protein is associated with height and flowering time (Lawit *et al*., 2010) and strongly associated with climate adaptation (Camus-Kulandaivelu *et al*., 2006, page). The rice *GHD7* locus has a strong effect on number of days to heading, number of grains per panicle, plant height and stem growth (Xue *et al*., 2008). In wheat, the dwarf gene *Rht-12* was shown to have a delaying effect on heading (Worland *et al*., 1994; Chen *et al*., 2013) as well as an additive interaction effect with *Ppd-D1* on plant height (Chen *et al*., 2018). Furthermore, it was shown that the tall *Rht-D1a* and the photoperiod sensitive *Ppd-D1b* allele positively affect leaf area and spike length throughout SE (Guo *et al*., 2018). To further examine the relationship among the different traits, we consider the following GWAS analysis using stringent correction of population structure.

The GWAS results indicate an independent genetic control of FH, temperature response and the timing of critical stages. Whereas vigour and FH as well as heading time, start and end of SE appear to be partly linked. Yet, FH could be predicted with surprising accuracy using the QTL for temperature response, vigour, start and end of SE which reflects the correlations found in the phenotypic data.

Previous studies investigating the control of developmental key stages in wheat with respect to temperature generally adopted the concept, that after fulfilment of photoperiod and vernalisation, *Eps* genes act as fine tuning factors independent of environmental stimuli (Kamran *et al*., 2014; Zikhali and Griffiths, 2015). Increasing temperature, apart from vernalisation is thought to generally quicken growth and development independent of the cultivar (Slafer and Rawson, 1995b; Porter and Gawith, 1999; Slafer *et al*., 2015). A genotype-specific temperature effect on the duration of different phases was not considered (Takahashi & Yasuda 1971, Slafer & Rawson 1995c). It was however reported, that photoperiod effects vary depending on temperature (Slafer and Rawson, 1995c). Under long days, Hemming et al. (2012) reported faster development and fewer fertile florets under high compared to low temperatures. Temperature-dependent effects were also found for different *Eps* QTL (Slafer and Rawson, 1995c; Gororo *et al*., 2001). It has previously been suggested, that *Eps* effects could be associated with interaction effects between genotype and temperature fluctuations (Slafer and Rawson, 1995c; van Beem *et al*., 2005).

The mechanisms of ambient temperature sensing and of its effects on growth and development are not yet well understood (Sanchez-Bermejo and Balasubramanian, 2016). However, important findings regarding ambient temperature effects on flowering time as well as on hypocotyl elongation have come from *Arabidopsis thaliana* (Wigge, 2013). With respect to these two traits, Sanchez-Bermejo and Balasubramanian (2016) reported distinct genotypic differences in temperature-sensitivity. According to their results, the flowering pathway genes *FRIGIDA* (*FRI*), *FLOWERING LOCUS C* (*FLC*) and *FLOWERING LOCUS T* (*FT*) are major candidate genes for ambient temperature mediated differences in flowering time (Sanchez-Bermejo and Balasubramanian, 2016). In the present study, we found *FRI* homologues near two of the three QTL for temperature response. *FRI* and *FLC* act as main vernalisation genes in *A. thaliana* (Johanson *et al*., 2000; Amasino and Michaels, 2010). In wheat, these genes are not yet well described. However, *FLC* orthologues were found to act as flowering repressors regulated by vernalisation in monocots (Sharma *et al*., 2017).

The most promising candidate gene for temperature response found near the QTL on chromosome 1B is *EARLY FLOWERING 3* (*ELF3*). In *Arabidopsis, ELF3* was found to be a core part of circadian clock involved in ambient temperature response (Thines and Harmon, 2010). In Barley, *ELF3* was shown to be involved in the control of temperature dependent expression of flowering time genes (Ejaz and von Korff, 2017). A mutant *ELF3* accelerated floral development under high ambient temperatures while maintaining the number of seeds (Ejaz and von Korff, 2017). Furthermore, *ELF3* has been reported as a candidate gene for *Eps1* in Triticum monococcum (Alvarez *et al*., 2016) as well as in wheat (Zikhali *et al*., 2016). A recent study in wheat showed an interaction between *Eps-D1* and ambient temperature which corresponded to different expression of *ELF3* (Ochagavía *et al*., 2019). In this study, we directly measured growth response to temperature during SE and found a significant MTA near *ELF3* on chromosome 1B. Following Ochagavía *et al*. (2019), this indicates that growth response to temperature is connected to *Eps-B1* which is homologue to *Eps-D1*. Furthermore, the reported temperature by *Eps-D1* interaction effects on heading by Ochagavía *et al*. (2019) are in agreement with the correlations found among growth response to temperature, GDD_15_, GDD_95_ and heading in the present study.

One important aspect we could not address in this study is the interaction of genotype specific temperature response with vernalisation and photoperiod (Slafer and Rawson, 1995c; Gol *et al*., 2017; Kiss *et al*., 2017). Due to the climate conditions in Switzerland, we expect fulfilment of vernalisation requirement in all genotypes. However, due to the broad geographic origin of the investigated genotypes, the relationship between temperature response and the timing of SE might be confounded by different photoperiod requirements. Nevertheless, the correlation between earliness and temperature response are in agreement with Ochagavia et al. (2019). It also remains unclear, if and to which extent temperature response varies across different developmental phases and how temperature response relates to other environmental stimuli such as vapour pressure deficit or radiation. Nevertheless, the results of this study present valuable information towards a better understanding of temperature response in wheat and may be of great importance for breeding. Temperature response could provide a breeding avenue for local adaptation as well as the control of plant height.

With the recent advancements in UAV-based phenotyping techniques, the growth of canopy cover and canopy height can be measured using image segmentation and structure from motion approaches (Bareth *et al*., 2016; Aasen and Bareth, 2018; Roth *et al*., 2018). Thus, temperature response can be investigated during the vegetative canopy cover development (Grieder *et al*., 2015) and during the generative height development as demonstrated here. It can also be assessed in indoor platforms (e.g. Parent and Tardieu, 2012) and the field using leaf length tracker (Nagelmüller *et al*., 2016) measuring short-term responses of leaf growth to diurnal changes in temperature. Combining this information may greatly improve our understanding about the genetic variation in growth response to temperature.

Together, the results of this study indicate that temperature response may be exploitable as a breeding trait to adjust phenology towards specific environments, either through phenotypic or marker assisted selection. Furthermore, a better understanding of temperature response may enhance the capability of crop models to predict crop performance under future climate change scenarios.

## Conclusion and outlook

Modern phenotyping platforms hold great promise to map the genetic factors driving the response of developmental processes to environmental stimuli. To the best of our knowledge, this is the first experiment dissecting the SE process into its underlying components: temperature-dependent elongation, temperature-independent vigour and elongation duration. The independent loci detected for these traits suggest, that it is possible to select them independently. The detected loci may be used to fine tune height and the beginning and end of SE as they explain a substantial part of the overall genotypic variation. With increases in automation, growth processes may be monitored in the field on a daily basis or even multiple times per day. This will increase the precision in assessing genotype responses to the fluctuation in meteorological conditions and will allow quantifying the relationship of these responses to yield. Remote sensing by means of unmanned aerial vehicles in combination with photogrammetric algorithms will allow to measure these traits in breeding nurseries. We believe that this is paving the road for a more informed selection to climate adaptation within individual growing seasons.

## Supplementary Data

**Fig. S1:** Correction of canopy height for spatial as well as random row and range effects.

**Fig. S2:** Summary of plot based linear model fits of stem elongation rate vs. temperature.

**Fig. S3:** Pearson correlation coefficients among 3-year BLUES of all investigated traits.

**Fig. S4:** Principal component analysis among marker genotypes.

**Fig. S5:** Manhattan plots and quantile-quantile plots depicting the GWAS results using the MLM approach.

**Fig. S6:** Manhattan plots and quantile-quantile plots depicting the GWAS results using the GLM approach.

**Table S1:** Genes of interest related to floral transition and flowering.

**Table S2:** Chromosome wise distance thresholds for LD-decay < r^2^ = 0.2

**Table S3:** Corresponding marker-trait associations for final canopy height with respect to Zanke et al. 2016.

**Table S4:** Selected putative candidate genes for headingGDD from the IWGSC reference genome annotation.

**Table S5:** Selected putative candidate genes for GDD_95_ from the IWGSC reference genome annotation.

**Table S6:** Selected putative candidate genes for GDD_15_ from the IWGSC reference genome annotation.

**Table S7:** 3-year BUEs of the investigated traits FH, headingGDD, GDD_15_, GDD_95_, GDD_SE_, time_SE_, slp_SER^~^T_, int_SER^~^T_.

## Data Availability

Processed phenotypic data is available as supplementary data. Unprocessed data and analysis scripts are available from the authors upon reasonable request.

## Acknowledgements

We sincerely thank Hansueli Zellweger for managing and nursing our field experiments. We further thank the members of the ETH crop science and the ETH molecular plant breeding groups, especially Michelle Nay and Beat Keller, for many fruitful discussions. We also thank Martina Binder for doing the correlation analysis between the FH data of this study and the data made by Zanke et al. (2014b) in the framework of her MSc-Thesis. We would like to thank Marion Röder (IPK Gatersleben) for supply of the GABI wheat panel including genetic information. Finally, we thank the anonymous reviewers for their helpful comments and suggestions. This work was supported by the Swiss National Foundation (SNF) in the framework of the project PhenoCOOL (project no. 169542).

## Author contribution

LK conducted the laser scans, did all statistical analyses and drafted the manuscript; SY assisted with the GWAS and candidate gene evaluation; MB assisted with the spatial correction; NK developed the analysis pipeline for the laser scans; AW drafted the grant application and supervised the overall concept; AH made the experimental design, developed the phenotyping models and assisted with the statistical analysis. All authors contributed to the drafting of the manuscript.

