## Supplementary Figures S1-S6 and Supplementary Tables S1-S3 for "Temperature response of wheat affects final height and the timing of stem elongation under field conditions"

**Fig. S1:** Correction of canopy height for spatial as well as random row and range effects.

**Fig. S2:** Summary of plot based linear model fits of stem elongation rate vs. temperature.

**Fig. S3:** Pearson correlation coefficients among 3-year BLUES of all investigated traits.

**Fig. S4:** Principal component analysis among marker genotypes.

**Fig. S5:** Manhattan plots and quantile-quantile plots depicting the GWAS results using the MLM approach.

**Fig. S6:** Manhattan plots and quantile-quantile plots depicting the GWAS results using the GLM approach.

**Table S1:** Genes of interest related to floral transition and flowering.

**Table S2:** Chromosome wise distance thresholds for LD-decay < $r^2$ = 0.2

**Table S3:** Corresponding marker-trait associations for final canopy height with respect to Zanke et al. 2016.

*See supplementary file 2 for tables S4 – S7:*

**Table S4:** Selected putative candidate genes for $heading_{GDD}$ from the IWGSC reference genome annotation.

**Table S5:** Selected putative candidate genes for $GDD_{95}$ from the IWGSC reference genome annotation.

**Table S6:** Selected putative candidate genes for $GDD_{15}$ from the IWGSC reference genome annotation.

**Table S7:** 3-year BUEs of the investigated traits FH, $heading_{GDD}$, $GDD_{15}$, $GDD_{95}$, $GDD_{SE}$, $time_{SE}$, $slp_{SER\sim T}$, $int_{SER\sim T.}$

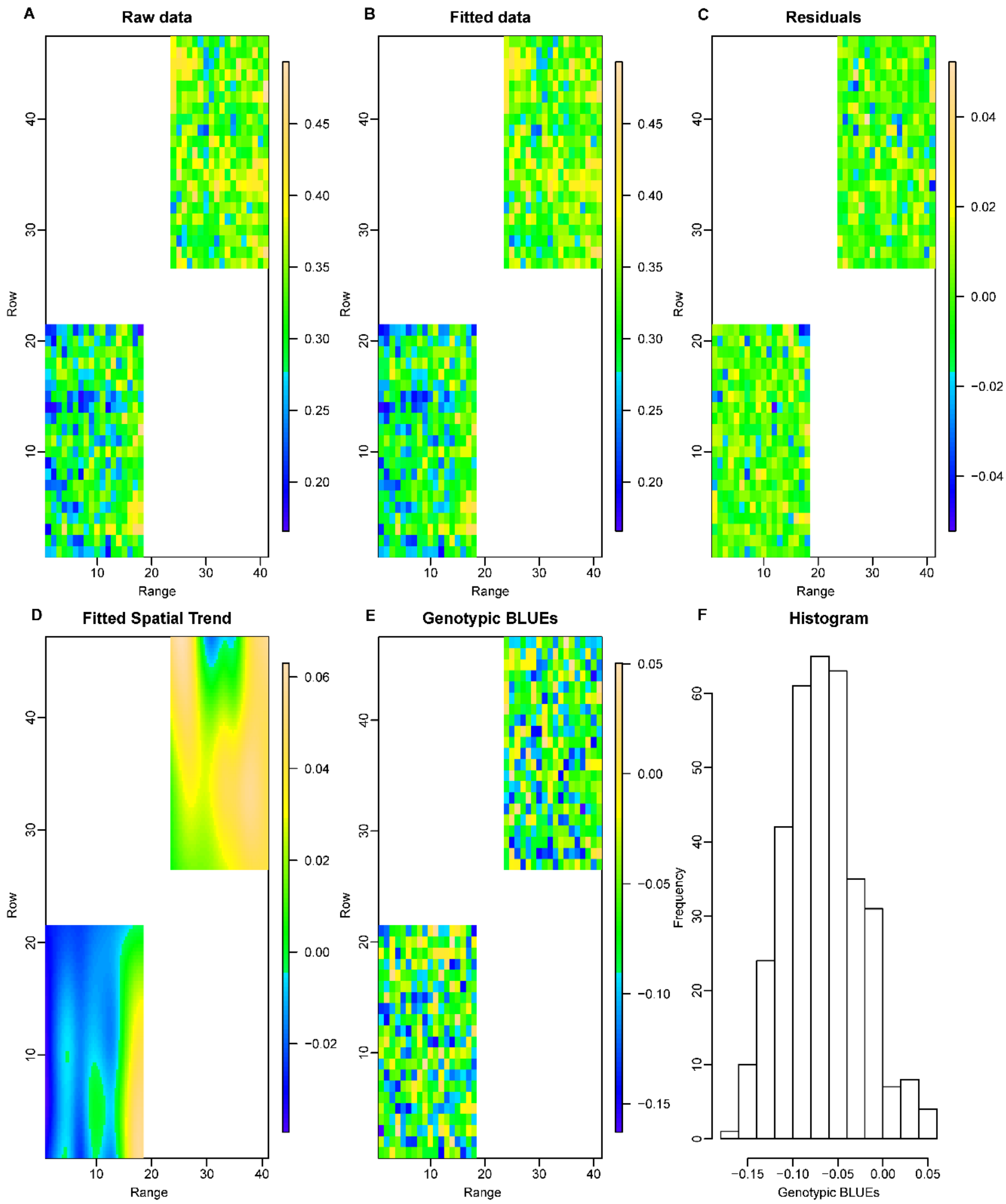


**Fig. S1: Correction of canopy height for spatial as well as random row and range effects.** The spatial correction of canopy height was done at each measurement time point by fitting a smoothed bivariate surface defined over rows and ranges to the raw canopy height data (see eq. 3 in Material and Methods) using the R-package SpATS (Rodríguez-Álvarez et al., 2018, Spatial Statistics). The figure shows the spatial correction for the time point 2017-05-09 displaying the raw canopy height data (A), the canopy height data fitted by the model (B), the residuals (C), the fitted spatial trend (D) as well as the genotypic estimates (E) and their distribution (F).

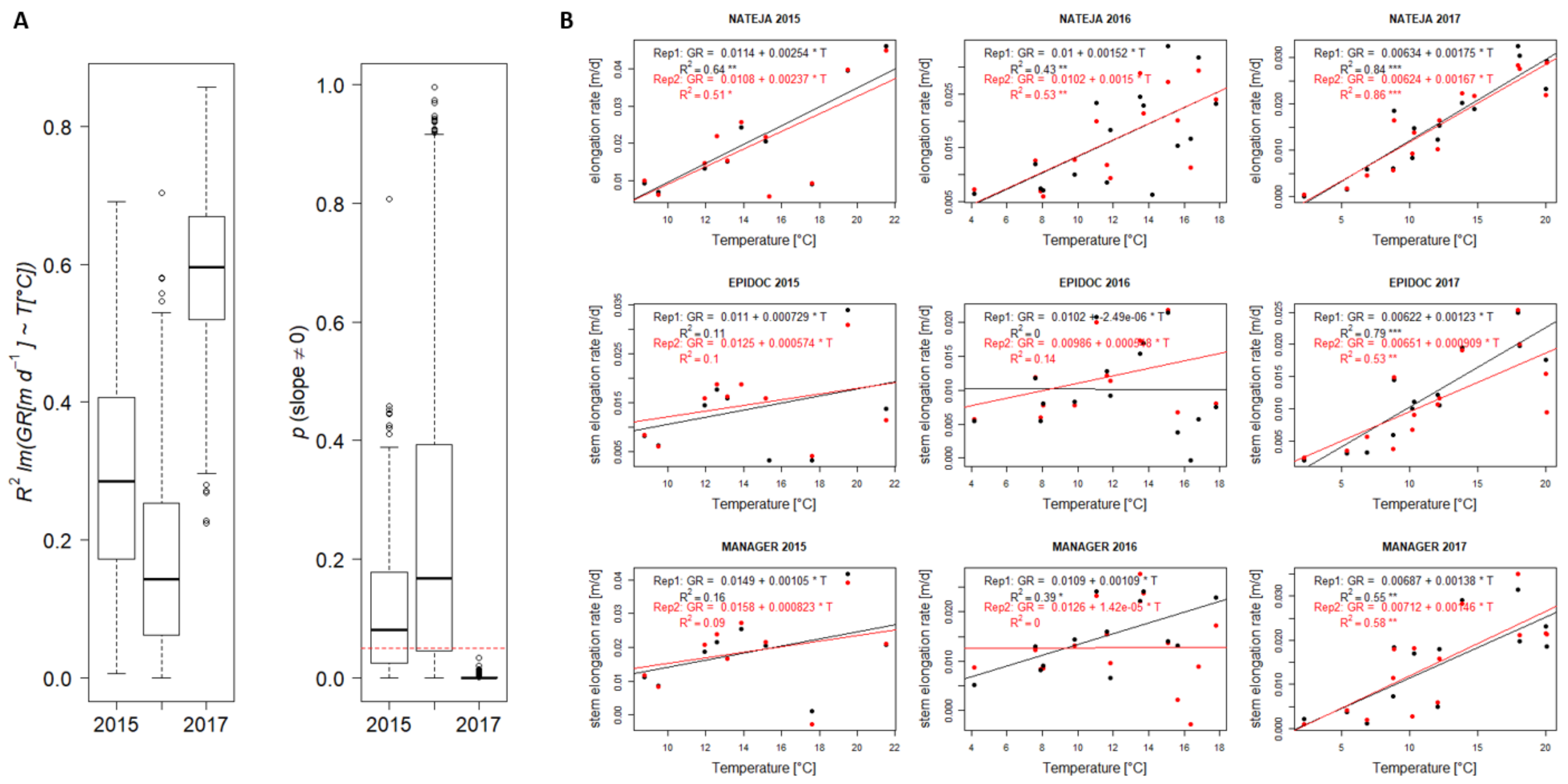


**Fig. S2: Summary of plot based linear model fits of stem elongation rate vs. temperature. A** Distribution of linear model $R^2$ and p-values of plot based linear model fits grouped by year. **B.** Genotype showing the best (NATEJA 2017 Rep1) and the two genotypes showing worst (EPIDOC 2016 Rep1; MANAGER 2016 Rep2) plot based linear model fit out of all plot based linear models fitted.

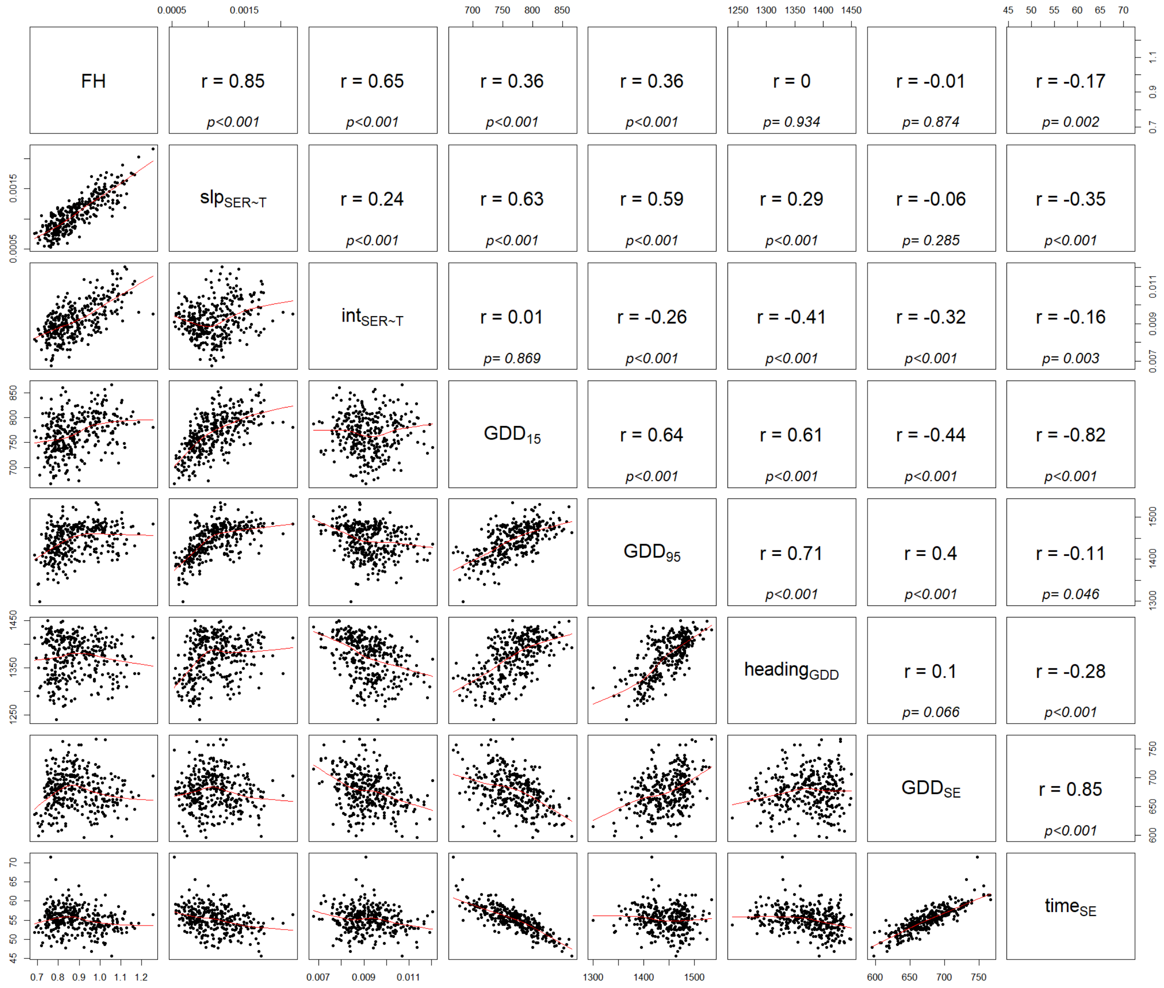


**Fig. S3: Pearson correlation coefficients among 3-year BLUEs of all investigated traits.**

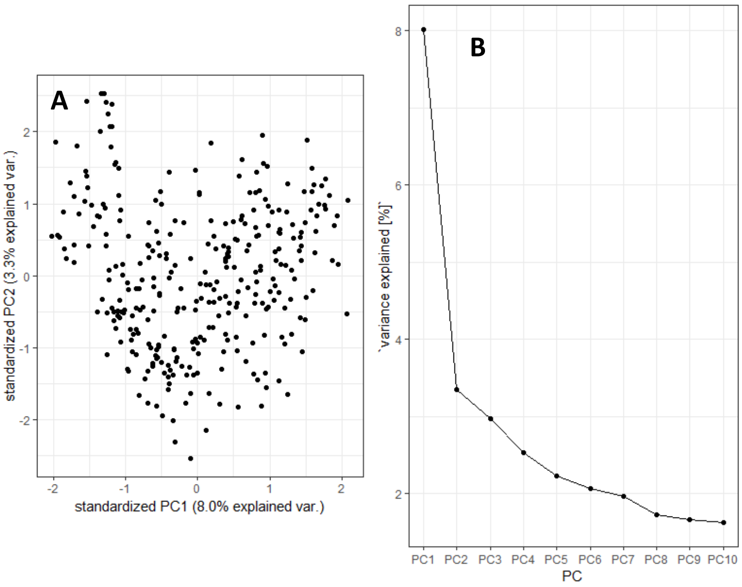


**Fig. S4: Principal component analysis among marker genotypes.** A: Biplot of the first two principal components. B: Screeplot showing the explained proportion of the variance for individual components.

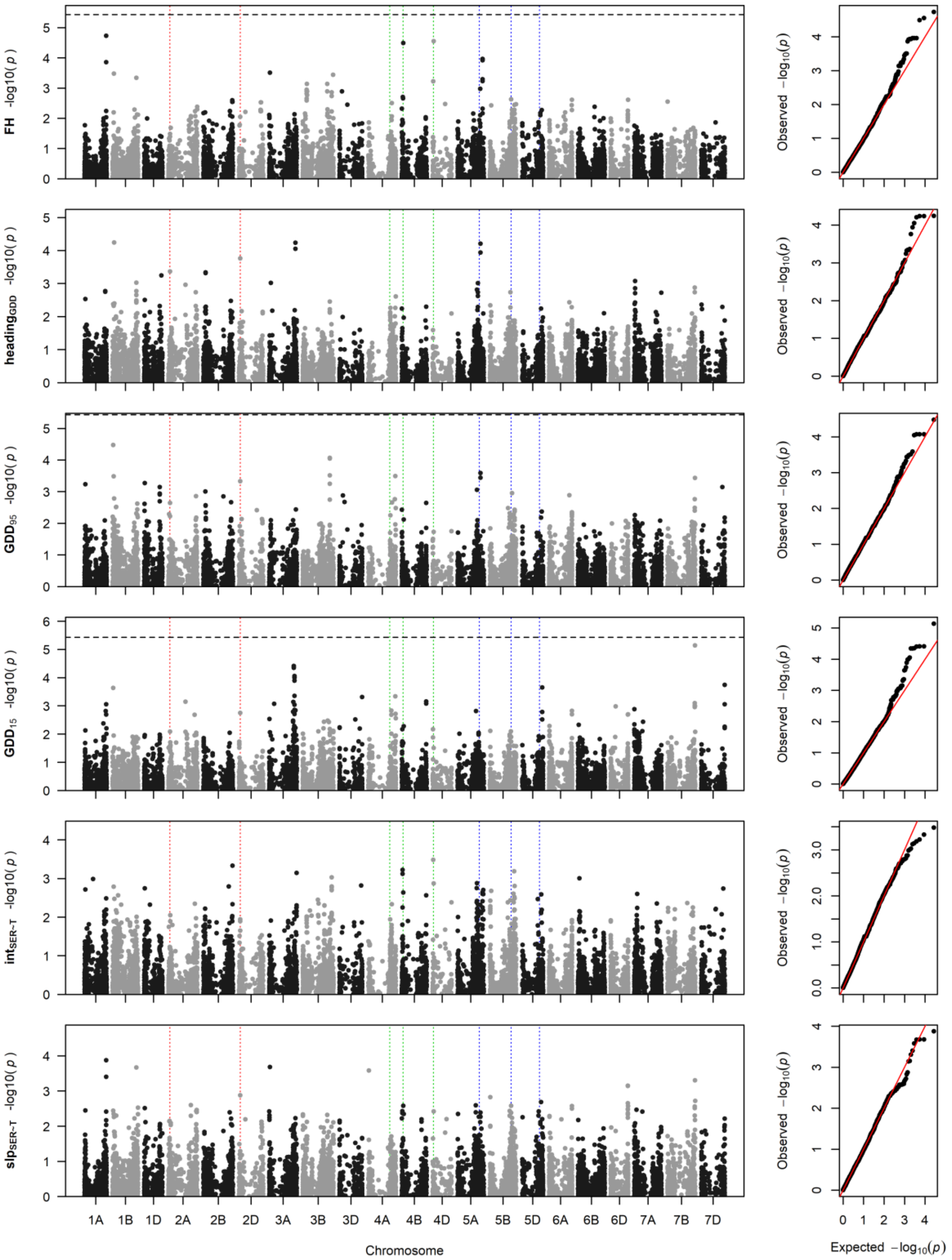


**Fig. S5: Manhattan plots and quantile-quantile plots depicting the GWAS results using the MLM approach** for final height (FH), heading in growing degree days ($heading_{GDD}$), GDD until start ($GDD_{15}$) and end ($GDD_{95}$) of stem elongation; vigour-related intercept ($int_{SER\sim T}$) and temperature-related slope ($slp_{SER\sim T}$) of stem elongation in response to temperature. Horizontal lines mark the Bonferroni corrected significance threshold for $p < 0.05$ (dashed line) and $p < 0.001$ (solid line). Vertical dotted lines mark the positions *Ppd-1* on chromosomes 2A and 2D (red), *Rht-1* on chromosomes 4A-4D (green) and *Vrn-1* on chromosomes 5A-5D.

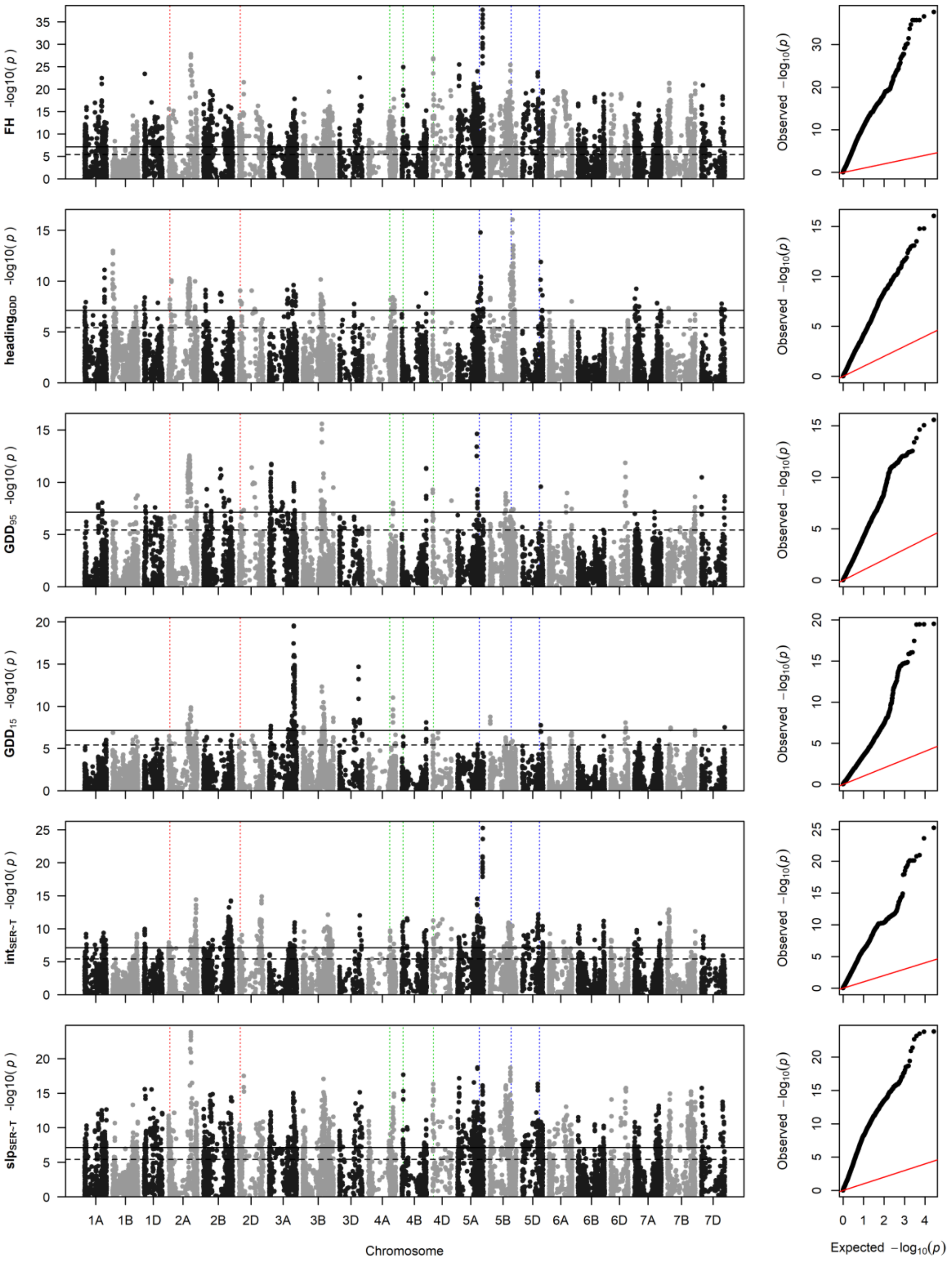


**Fig. S6: Manhattan plots and quantile-quantile plots depicting the GWAS results using the GLM approach** for final height (FH), heading in growing degree days (heading$_{GDD}$), GDD until start (GDD$_{15}$) and end (GDD$_{95}$) of stem elongation; vigour-related intercept (int$_{SER\sim T}$) and temperature-related slope (slp$_{SER\sim T}$) of stem elongation in response to temperature. Horizontal lines mark the Bonferroni corrected significance threshold for $p < 0.05$ (dashed line) and $p < 0.001$ (solid line). Vertical dotted lines mark the positions *Ppd-1* on chromosomes 2A and 2D (red), *Rht-1* on chromosomes 4A-4D (green) and *Vrn-1* on chromosomes 5A-5D.

**Table S1:** Genes of interest related to floral transition and flowering. Genome positions (r.start-r.end) were derived by blasting the respective published sequence (GenBank ID; https://www.ncbi.nlm.nih.gov/genbank/) against the IWGSC reference genome.

| Gene | Chr | r.start | r.end | GenBank ID |
|---|---|---|---|---|
| *Ppd-A1* | chr2A | 36'934'562 | 36'933'892 | DQ885753.1 |
| *Ppd-D1* | chr2D | 33'953'359 | 33'952'698 | DQ885766.1 |
| *Rht-A1* | chr4A | 582'479'578 | 582'477'716 | JF930277.1 |
| *Rht-B1* | chr4B | 30'861'382 | 30'863'247 | JX993610.1 |
| *Rht-D1* | chr4D | 18'781'062 | 18'782'933 | AJ242531.1 |
| *TaELF3* | chr1D | 493'485'605 | 493'484'553 | KR055809.1 |
| *TaFT3-A1* | chr1A | 528'066'476 | 528'066'282 | KX161737.1 |
| *TaFT3-B1* | chr1B | 581'414'952 | 581'414'758 | KX161739.1 |
| *TaFT3-D1* | chr1D | 430'469'335 | 430'469'144 | KX161740.1 |
| *Vrn-A1* | chr5A | 587'423'240 | 587'423'056 | AY616452.1 |
| *Vrn-B1* | chr5B | 573'815'903 | 573'815'719 | AY747603.1 |
| *Vrn-B3* | chr7B | 9'703'464 | 9'703'735 | DQ890162.1 |
| *Vrn-D1* | chr5D | 467'184'278 | 467'184'094 | AY747597.1 |
| *Vrn-D4* | chr5D | 467'184'278 | 467'184'094 | KR422424.1 |

**Table S2** Chromosome wise distance thresholds for LD-decay < $r^2$ = 0.2

| Chromosome | $r^2$ threshold [bp] |
|---|---|
| 1A | 6'161'631 |
| 1B | 17'286'788 |
| 1D | 12'822'505 |
| 2A | 10'381'104 |
| 2B | 11'361'169 |
| 2D | 10'416'001 |
| 3A | 9'887'297 |
| 3B | 5'969'718 |
| 3D | 4'233'196 |
| 4A | 4'298'677 |
| 4B | 11'488'268 |
| 4D | 4'220'358 |
| 5A | 10'344'140 |
| 5B | 17'533'559 |
| 5D | 4'862'712 |
| 6A | 4'692'731 |
| 6B | 19'679'489 |
| 6D | 1'416'632 |
| 7A | 7'133'812 |
| 7B | 6'604'947 |
| 7D | NA |

**Table S3** Corresponding marker-trait associations for final canopy height with respect to Zanke et al. 2014*b*. MTA Zanke et al 2014*b* denotes marker trait associations reported by Zanke et al. 2014*b*, Closest MTA denotes the closest respective associated marker found in this study. Distance gives the distance in base pairs and $r^2$ the pairwise linkage disequilibrium between the two respective SNP.

| Chr | MTA Zanke et al. 2014*b* | Closest MTA | distance | $r^2$ |
|---|---|---|---|---|
| 2B | Excalibur_c23185_155 | wsnp_Ku_c11665_18999583 | 1'679'482 | 0.25 |
| 5A | wsnp_Ex_c23795_33033959 | wsnp_Ku_rep_c71232_70948744 | 1'457 | 0.99 |
| 5A | Kukri_c75091_220 | wsnp_Ku_rep_c71232_70948744 | -5'660 | 0.99 |
| 5B | wsnp_Ex_c5155_9140608 | BS00109560_51 | 2'442'796 | 0.82 |
| 5B | wsnp_Ex_c54092_57099525 | BS00109560_51 | -4'850'924 | 0.55 |
| 6A | BS00062823_51 | BS00022120_51 | 51'119'779 | 0.60 |
